# An Automated High-throughput Affinity Capture-Mass Spectrometry Platform with Data- Independent Acquisition

**DOI:** 10.1101/2024.08.13.607785

**Authors:** Hui Jing, Paul L. Richardson, Gregory K. Potts, Sameera Senaweera, Violeta L Marin, Ryan McClure, Adam Banlasan, Hua Tang, James E. Kath, Renze Ma, Jon D. Williams

**Author notes:** Contributed equally (Hui Jing, Paul L. Richardson).

## Abstract

Affinity capture (AC) combined with mass spectrometry (MS)-based proteomics is highly utilized throughout the drug discovery pipeline to determine small molecule target selectivity and engagement. However, the tedious sample preparation steps and time-consuming MS acquisition process has limited its use in high-throughput format. Here, we report an automated workflow employing biotinylated probes and streptavidin magnetic beads for small molecule target enrichment in 96-well plate format, ending with direct sampling from EvoSep Solid Phase Extraction tips for liquid chromatography (LC)-tandem mass spectrometry (MS/MS) analysis. The streamlined process significantly reduced both overall and hands-on time needed for sample preparation. Additionally, we developed a data-independent acquisition-mass spectrometry (DIA-MS) method to establish an efficient label-free quantitative chemical proteomic kinome profiling workflow. DIA-MS yielded coverage of ∼380 kinases, a >60% increase compared to using a data-dependent acquisition (DDA)-MS method and provided reproducible target profiling of the kinase inhibitor dasatinib. We further showcased the applicability of this AC-MS workflow for assessing the selectivity of two clinical-stage CDK9 inhibitors against ∼250 probe-enriched kinases. Our study here provides a roadmap for efficient target engagement and selectivity profiling in native cell or tissue lysates using AC-MS.

## Introduction

Mass Spectrometry (MS)-based chemical proteomics coupled with affinity-^1^ or activity-based^2^ protein capture allows simultaneous assessment of the selectivity and potency of small molecule inhibitors against their natively expressed targets in cells or tissue. Typical affinity capture (AC) experiment utilizes probe compounds that are immobilized on a solid support to enrich target proteins from cell or tissue lysates. Using quantitative MS as a readout, the target profile for a small molecule inhibitor can be determined by competition experiments measuring the reduction of target proteins enriched by the immobilized probe matrix as a function of the free inhibitor concentration^3^.

Chemical probes with broad inhibition profiles within a target class can be combined and used to enrich a large fraction of a protein family’s members, thereby enabling a binding assay for an entire target class. For kinases, AC-MS-based profiling was previously developed by combining several agarose bead-immobilized nonselective kinase inhibitors^3^. Current state-of-the-art methodologies allow the assessment of kinase inhibitor selectivity against >250 quantified kinases in one experiment^4-9^. Although the workflow has been progressively miniaturized^6, 8, 9^ and improved for a higher throughput^6, 8^, sample preparation bottlenecks remain. For example, enrichment with probe-coated beads requires centrifugation^8, 9^ or filtration^6^ to remove the unbound proteome and exchange sample buffer. Further, filter plate-assisted wash steps are often followed with elution of bead-bound proteins and SDS-PAGE for subsequent in-gel digestion^4, 5^, which makes processing a large number of samples challenging.

In considering ways to streamline and automate the AC-MS process, we looked to combine biotinylated chemical probes and streptavidin magnetic (SA-Mag) beads to allow efficient and robot-assisted enrichment, washing and proteolytic digestion in one pot. Although streptavidin- or other avidin-coated magnetic beads have been used with biotinylated small molecules^10^, nucleic acids^11-13^ and proteins^14^ to capture and detect interacting proteins, SA-Mag beads suffer from challenges to pull down certain target proteins^15^ and produces intense streptavidin contamination tryptic peptides that impedes MS-based target protein identification^16^. For these reasons, SA-Mag beads combined with biotinylated chemical probes have not yet been effectively used to enrich hundreds of members from a target protein class, such as kinases, unlike the widely used agarose bead-immobilized probes, which are not as amenable to automation.

Additionally, state-of-the-art AC-MS chemoproteomic workflow typically employs liquid chromatography (LC) times ranging from one to three hours per sample^4-9^, which limits its application in drug discovery. Despite the significant improvements in multiplexing up to 18 samples in a single MS run through isobaric labeling^6, 17^, the labeling reagents are expensive and the cross-comparison of larger number of samples is challenging. On the other hand, label free proteomics is relatively cost- and time-effective, and more flexible for comparisons among a large set of samples. With advanced data processing software, data independent acquisition (DIA)^18^ methods have enabled comprehensive proteome profiling from complex samples with short gradients due to their high coverage and excellent quantitative performance^19-21^.

In this study, we present the step-by-step development of a streamlined and automated SA-Mag bead-based AC-MS workflow. This workflow was optimized in a 96-well format with minimal overall and hands-on time and utilizes DIA-MS analysis for efficient chemoproteomic kinome profiling, resulting in a coverage of approximately 400 kinases and robust quantification of target engagement against ∼250 probe-enriched kinases. Our results demonstrate a highly streamlined one-pot AC-MS workflow yielding comprehensive coverage with high-throughput amenable LC gradient and low sample input. Moreover, we applied this platform to profile two clinical stage CDK9 inhibitors, showcasing its effectiveness to selectivity evaluation of drug candidates.

## Experimental Section

### Chemical synthesis

Synthesis and characterization of chemical probes are described in Supporting Information 1.

### Cell culture

K562 (chronic myeloid leukemia), THP-1 (acute monocytic leukemia) and NCI-H1155 (non-small cell lung cancer) cells were originally obtained from American Type Culture Collection (ATCC). MV4-11 (acute monocytic leukemia) cells were obtained from Leibniz Institute DSMZ. Cells were cultured in RPMI-1640 medium (11875093, Gibco) supplemented with 10% fetal bovine serum (FBS, A3840202, Gibco), penicillin (100 units/mL) and streptomycin (100 μg/mL). Upon reaching ∼80% confluence, cells were harvested by centrifuging at 1,400 g for 3 min at 4 °C. Cell pellets were washed 2 times using 1x ice-cold DPBS before being flash frozen and stored at -80 °C until further processing.

### Cell lysis

K-562, THP-1, MV4-11, and NCI-H1155 cells were lysed in lysis buffer (0.5% Triton X-100, 50 mM HEPES pH 7.5, 150 mM NaCl), supplemented with 1.5 mM MgCl_2_, 250 U/mL of Benzonase (E1014, EMD Millipore), and protease inhibitors cocktail (04693132001, Roche) by incubating on ice for 30 min with occasional vortexing and centrifuging at 4 °C, 21,000 g for 30 min. Supernatant was transferred to new tubes and protein concentration was determined by Pierce^™^ 660nm assay (22662, Thermo Fisher Scientific) and adjusted to 2.5 mg/mL. When mixed cell lysates were used, the lysate from each cell line was mixed at a 1:1:1:1 ratio for downstream processing.

### Affinity capture using Cp19-Affi-Gel^™^ 10

Cp19-Affi-Gel^™^ 10 (3.5 nmol effective Cp19/sample) stored in isopropanol was washed 2 times with water and 3 times with lysis buffer by centrifuging the beads in 1.5 mL tube at 2,000 g for 1 min. After the last wash, lysis buffer was removed by pipetting, 200 µL of K562 cell lysates (2.5 mg/mL) was added to the beads and incubated at 4 °C on thermomixer (5382000023, Eppendorf) at 1,000 rpm for 2 hours. The beads were separated from lysate supernatant by centrifugation at 2,000 g for 1 min, then washed in lysis buffer (400 µL/well) for 4 times, followed by wash buffer (50 mM HEPES pH 7.5, 0.2% deoxycholate) for another 4 times. Bound proteins were processed for western blot or MS-based proteomic analysis as described below.

### Magnetic beads-based affinity capture

Pierce^™^ streptavidin magnetic beads (SA-Mag, 10 mg/mL, 25-200 μL slurry/sample, 88816, Thermo Fisher Scientific) were prewashed 3 times with 400 μL of lysis buffer and resuspended in 200 μL lysis buffer in 1.5 mL tube or each well of 96-deepwell plate (951032603, Eppendorf). SA-Mag beads were separated from supernatant using DynaMag^™^-2 magnet (12321D, Thermo Fisher Scientific) for the manual workflow or 96-well magnetic-ring stand (AM10050, Thermo Fisher Scientific) for the automated workflow. DMSO, Cp19-biotin probe or biotinylated kinome probes (mixed at 1:1:1:1:1) were spiked in the resuspended SA-Mag beads at final concentration of 17.5 µM. Beads were incubated with probes at 4 °C on thermomixer at 1,000 rpm overnight, washed 3 times using lysis buffer (400 μL/sample) and placed on ice until cell lysates were ready for pull-down.

For initial method optimization where compound was not tested, 200 µL/sample of cell lysate was incubated with prepared probe-bound beads at 4 °C on thermomixer at 1,000 rpm for 2 hours. When kinase inhibitor was tested, 200 µL/sample of cell lysate was incubated with dasatinib, BAY-1143572 or BAY-1251152 at indicated final concentrations, and incubated at 4 °C on thermomixer at 1,000 rpm for 45 min. Compound-treated lysates were then incubated with probe-bound beads for 30 min at 4 °C. For the tandem pull-down, supernatant from DMSO-treated cell lysate was collected after the incubation with beads and subjected to another 30 min incubation with fresh probe-bound beads. The resulting beads from above were washed in lysis buffer (400 µL/well) for 4 times, followed by wash buffer (50 mM HEPES pH 7.5, 0.2% deoxycholate) for another 4 times. Bound proteins were processed for western blot or MS-based proteomic analysis as described below.

### Immunoblot

Proteins pulled down on beads were resuspended in 40 µL of 2x LDS sample buffer (NP0007, Thermo Fisher Scientific) and heated at 70 °C for 10 min. Proteins from 10 µL of supernatant were resolved on NuPAGE^™^ 4-12% Bis-Tris mini protein gel (NP0321BOX, Thermo Fisher Scientific) and transferred using iBlot^T™^ 2 dry blot system (Thermo Fisher Scientific) by following manufacturer’s instructions. The membrane was blocked with Intercept^®^ (TBS) blocking buffer (927-60001, Licor) at room temperature for 1 hour and incubated with primary antibody BTK mAb (1:1000, 8547, Cell Signaling) in Intercept^®^ T20 (TBS) antibody diluent (927-65001, Licor) overnight at 4 °C, followed by IRDye^®^ 800CW Goat anti-Rabbit IgG secondary antibody (1:5000, 926-32211, Licor) at room temperature for 1 hour. After several washes with TBST, images were acquired by ODYSSEY^®^ DLx imaging system (Licor).

### On-bead digestion following affinity capture

Bead-bound proteins were processed using PreOmics iST kit (P.O.00027). Beads that were washed above were resuspended in 100 µL LYSE and heated at 60 °C in thermomixer at 1,000 rpm for 10 min. 50 µL DIGEST was added for trypsin digestion at 37 °C in thermomixer at 1,000 rpm for 1 hour. Digestion was quenched and peptides were cleaned up following manufacturer’s instruction. The resultant peptide samples were frozen, dried by vacuum centrifugation and stored at −80 °C until further analysis.

### LC–MS/MS

Dried peptides were resuspended in water with 0.1% formic acid and analyzed on the EvoSep One and Orbitrap Exploris^™^ 480 mass spectrometer. Half of the resuspended peptides and 0.2 µL of iRT (10x) peptides (1816351, Biognosys) were loaded onto Evotip (EV2001, EVOSEP) following manufacturer’s manual. The loaded peptides were separated on an EASY-Spray^™^ ES906 column (15 cm x 150 uM ID, 2uM particle size). using the EvoSep 30SPD 44 min gradient. The Exploris^™^ 480 was operated in data-dependent acquisition (DDA) or data-independent acquisition (DIA) manner as described below.

#### DDA-MS

The Exploris DDA method used survey MS resolving power of 120,000, with a full scan range of 300-1800 m/z. Survey MS scans were collected with an AGC of 300% (1.2eE6) and “custom” maximum injection times of 25 msec. The MS RF lens was set to 40%, and the total cycle time of the MS was set to 3 seconds per cycle. Advanced Peak Detection (APD) was utilized for all DDA runs, along with monoisotopic precursor selection (MIPS). Peptides were required to be fragmented after reaching a minimum intensity of 5000 with a charge state between +2-8. Dynamic exclusion times of 15 seconds were utilized after a peptide was fragmented, with exclusion tolerances of +/-10 ppm. MS/MS isolation windows of 1.2 m/z were utilized for fragmenting peptides by HCD using a normalized collision energy of 30%. All fragments were analyzed in the Orbitrap using a resolving power of 30,000. The MS/MS scan range was collected with “custom” maximum injection times of 75 msec and a standard MS/MS AGC target of 100% (50,000). All data was collected in centroid mode.

#### DIA-MS

The Exploris DIA method used survey MS resolving power of 60,000, with a full scan range of 350-1200 m/z. Survey MS scans were collected with an AGC of 100% (4E5) and maximum injection time to “Auto.” The MS RF lens was set to 40%, and the total cycle time of the MS was set to 3 seconds per cycle. Notably, custom DIA windows were designed using the method editor’s “targeted MS” (tMSn) settings to allow custom window sizes to be employed across the m/z range. Peptides were fragmented by HCD using a normalized collision energy of 30% and analyzed using Orbitrap with a resolving power of 30,000. The MS/MS scan range was collected from 145-1450 m/z, with “custom” maximum injection times of 54 msec and a MS/MS AGC target of 100% (50,000). All data was collected in centroid mode. 70 total DIA windows were utilized across the m/z range from 400 m/z to 1000 m/z, with each isolation window size spanning from between +/-5 m/z to 25 m/z (see Supporting Information 2).

### Peptides and protein identification and quantification

The raw mass spectrometry data was deposited in MassIVE (https://massive.ucsd.edu/ProteoSAFe/static/massive.jsp) with the data set identifier MSV000095587. Raw DDA-MS and DIA-MS data files were searched with MaxQuant software (v.2.0.1.0)^22^ and Spectronaut^™^ 17 software (Version 17.4.230317), respectively, using standard settings unless otherwise described. Tandem mass spectra were searched against all protein sequences as annotated in the UniProt human proteome reference database (UP000005640, with isoforms, 2021-01-04). Carbamidomethylated cysteine was set as fixed modification. Variable modifications included oxidation of methionine and N-terminal protein acetylation. Trypsin/P was specified as proteolytic enzyme with up to two missed cleavage sites. For DDA-MS, label-free quantification and match between runs were enabled. Results were filtered for 1% peptide and protein false discovery rate (FDR) using a target-decoy approach using reversed protein sequences. For DIA-MS, precursors were filtered using Q value cutoff.

### Data analysis of kinase pulldown and inhibition

The MaxQuant search file (proteinGroup.txt) and Spectronaut output file of DDA-MS and DIA-MS data, respectively, were used for subsequent filtering and analysis using Perseus (2.0.10.0). Unless otherwise indicated, reverse hits, potential contaminants and proteins identified in < 60% replicates of the Vehicle control kinase pull-down group and/or with one peptide were discarded. Protein intensity values – LFQ (label-free quantification) intensity and PG (Protein Group) Quantity for DDA- and DIA-MS, respectively – were used to obtain fold change for kinase enrichment and statistical analysis by Student’s t-tests (two sided) using log-transformed intensities. Statistical tests were corrected for multiple testing by an FDR of 5%. Kinases showing significant enrichment over the no probe group (fold change ≥ 2, *P* ≤ 0.05) were further analyzed for inhibition by compound. Protein intensities were normalized to the average DMSO control intensity to obtain relative residual binding intensities for each protein group at every inhibitor concentration. IC_50_ values were obtained for the proteins with significantly affected intensity at highest inhibitor concentration (*P* ≤ 0.05) from GraphPad Prism 10 using the log(inhibitor) vs. normalized response (variable slope) function. *K*_d_ values were calculated by multiplying IC_50_ values with a correction factor that accounts for kinase depletion from the lysate by immobilized probes. The depletion was measured by the ratio of the intensity of a kinase captured in the second over that of the first of the two consecutive pulldowns of the same DMSO control lysate.

## Results

### Automated AC-MS workflow development

To establish an automatable AC-MS workflow, we set out to test if a biotinylated chemical probe combined with SA-Mag beads enriches known protein targets. For this purpose, we selected compound 19 (Cp19), a potent and nonselective tyrosine kinase inhibitor, that has been reported to enrich over 200 kinases from cell lysates when immobilized on agarose beads^4, 23^. We reasoned that its broad kinome coverage would allow us to assess the effectiveness of the automated workflow. To compare with the well-established agarose affinity matrix-based approach, Affi-Gel^™^ 10-conjugated^24^ and biotinylated variants of Cp19 were synthesized, with the former containing a PEG2 linker, and the latter containing PEG4, 8 or 12 linkers (Fig. 1A). The initial method development experiments were performed with K562 cells, which have been previously used for kinome profiling^4, 5^.

**Fig 1.**
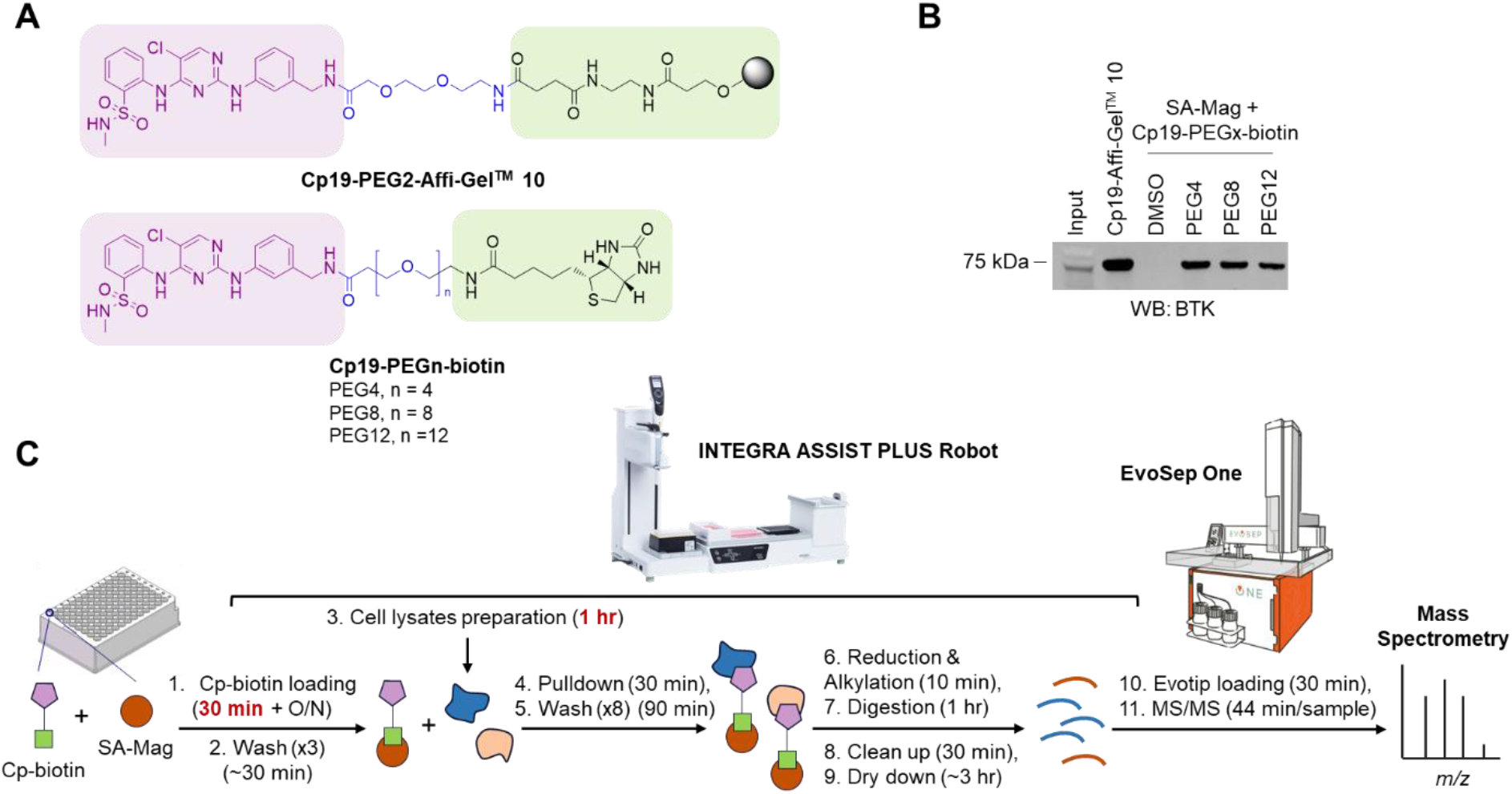
A magnetic bead-based automatable AC-MS chemoproteomics workflow. (A) Structures of Cp19-PEG2-Affi-Gel^™^ 10 and Cp19-PEG4/8/12-biotin. The nonselective kinase inhibitor Cp19 moiety is highlighted in purple, and the Affi-Gel^™^ 10 or biotin affinity moiety is highlighted in green. (B) Western blot (WB) detection of BTK enrichment by Cp19-PEG4/8/12-biotin using SA-Mag beads in comparison to that by of Cp19-PEG2-Affi-Gel^™^ 10 from K562 cell lysates. (C) Schematic overview of the automated AC-MS chemoproteomics workflow. The time that each step takes for a 96-well plate of samples is indicated. Highlighted in red is the time for the steps that mainly require manual preparation.

To test the pull-down efficiency of using biotinylated probes immobilized on SA-Mag beads, we pre-loaded 1 mg of SA-Mag beads with 3.5 nmol of the Cp19-biotin probe, corresponding to one equivalent of bead binding capacity. We then incubated the immobilized Cp19-biotin probe or Cp19-Affi-Gel^™^ 10 (3.5 nmol Cp19 effective concentration) with 200 μL of K562 cell lysates (2.5 mg/mL). Western blot analysis of affinity captured-proteins showed that the Cp19-biotin probes enriched Bruton’s tyrosine kinase (BTK), a known Cp19 target ^4^, albeit with lower efficiency compared to Cp19-Affi-Gel^™^ 10. We noted that as the PEG linker length increased, less BTK was pulled down by the biotinylated Cp19 probes, suggesting potential impact of linker length on target enrichment (Fig. 1B).

Having confirmed that Cp19-biotin probes immobilized on SA-Mag beads enrich a known Cp19 target, we next developed a 96-well plate-based automatable AC-MS sample preparation and analysis workflow. An INTEGRA Assist Plus pipetting robot was used to automate washing and MS sample preparation steps involving repetitive pipetting. The use of SA-Mag beads allowed us to use a magnet plate for easy separation of beads from the liquid phase, streamlining the process of probe immobilization, target enrichment, washing, and trypsin digestion all in one pot. To enable high-throughput MS sample acquisition, product peptides were loaded on Solid Phase Extraction (SPE) Evotips ^11^, also in a 96-well format, using the Assist Plus robot, and subsequently analyzed by LC-MS/MS using an EvoSep One LC system^12^ and Orbitrap Exploris 480 MS with a 44-minute gradient and label-free data-dependent acquisition (DDA)-MS method (Fig. 1C). Based on the automation protocol set up on the INTEGRA Assist Plus, we determined the time needed for each step (Fig. 1C) and the entire sample preparation process – for one 96-well plate of samples, approximately 1.5 days with one over-night incubation, one optional pause point and only ∼2 hours of hands-on time for probe and cell lysate preparation.

### Optimization of the automated AC-MS workflow

We next sought to optimize the automated workflow to maximize target capture and coverage by using Cp19-PEG8-biotin as a tool probe. As steric hindrance that results from multisite attachment of probe compounds could alter affinity agent’s activity^25, 26^, we first evaluated whether the ratio between SA-Mag and biotinylated probe could affect Cp19 target pull-down. For 0.5 mg of K562 cell lysate, a fixed 2 mg of SA-Mag beads (200 μL of 10 mg/mL bead slurry, binding capacity of ∼7 nmol biotinylated fluorescein) was pre-incubated with Cp19-PEG8-biotin at 0 – 2× equivalent bead binding capacity. Through DDA-MS analysis, the median and overall MS intensity (hereafter referred to as intensity) for quantified kinases increased as a function of the added Cp19-PEG8-biotin and reached a plateau at 7 – 14 nmol of Cp19-PEG8-biotin, corresponding to 1 – 2 × SA-Mag binding capacity (Fig. 2A, Table S1). Saturating SA-Mag beads with Cp19-PEG8-biotin did not compromise the intensity for kinases, suggesting that there was unlikely to be steric hindrance between the immobilized biotinylated probes that limits target capture. While kinase intensities were dependent on probe addition, number of kinases and non-kinase proteins detected were not affected by probe concentration (Fig. S1A), demonstrating background binding of kinases and non-kinases to SA-Mag beads.

**Fig 2.**
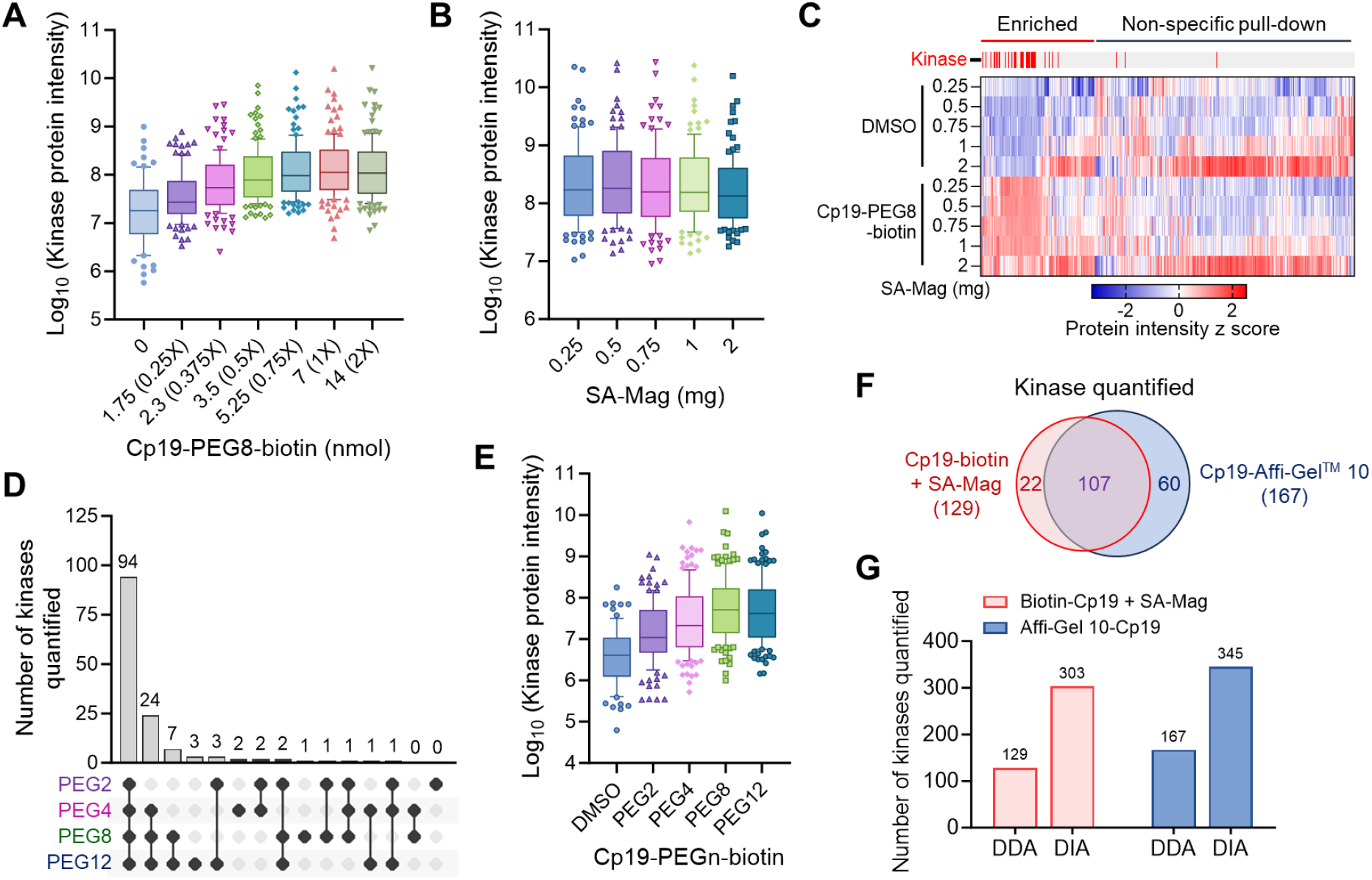
Optimization of the AC-MS workflow for chemoproteomics. (A, B) MS intensity distributions for kinases quantified by using (A) a fixed amount of SA-Mag beads (2 mg) and varying Cp19-PEG8-biotin loading or (B) the indicated amount of streptavidin magnetic beads (SA-Mag) and a fixed a probe/bead ratio of 3.5 nmol/mg. The x-axis in (A) lists both the absolute amount of Cp19-PEG8-biotin in nmol and, in parantheses, corresponding number of equivalents relative to the binding capacity of 2 mg SA-Mag (3.5 nmol/mg). (C) Heatmap showing the relative quantities of proteins identified in the experiment in (B). Columns corresponding to individual proteins are labeled above the heatmap in red for kinases and in grey for non-kinases. Protein intensity Z-score values were generated using log2 transformed protein intensity values in Perseus and plotted for each protein. (D) Upset plot showing the overlap of quantified kinases by Cp19-biotin probes with PEG2, 4, 8, and 12 linkers. Kinases quantified in at least two out of three replicates were plotted. (E) Intensity distributions for kinases quantified by using Cp19-biotin probes with PEG2, 4, 8 and 12 linkers. (F) Venn diagram illustrating the number of kinases quantified by the automated workflow using Cp19-PEG8-biotin in combination with SA-Mag and the manual workflow using Cp19-Affi-Gel^™^ 10. (G) Comparison of the number of kinases quantified between DDA-MS and DIA-MS for both the manual and automated workflows. All experiments were performed with three independent replicates per group. For (A), (B) and (E), Log_10_(kinase protein intensity) indicates median Log_10_(LFQ intensity) for each quantified kinase across replicates.

Boxplot shows the median central line and extends from the 25th to 75th percentiles. Whiskers represent protein quantity within the 10th to 90th percentiles.

We then evaluated the impact of the amount of SA-Mag beads on background binding and kinase enrichment while keeping Cp19-PEG8-biotin fixed at 1 × binding capacity. The intensity for quantified kinases was not significantly affected by titrating SA-Mag beads from 0.25 to 2 mg of beads, with 2 mg showing a modest negative impact (Fig. 2B, Table S2). As expected, most quantified kinases (97 out of 112) at any SA-Mag condition were ≥2-fold enriched when comparing experiments with and without Cp19 probe (Fig. 2C). While the number of kinases quantified went up modestly with more SA-Mag beads, all proteins quantified exhibited a greater increase (Fig. S1B), suggesting non-specific pull-down as a function of the amount of SA-Mag beads. Comparing the intensity of all quantified proteins with and without Cp19-PEG8-biotin indicated that more SA-Mag beads, especially at 2 mg, afforded greater intensity for those that were non-specifically pulled down (Fig. 2C). Based on these results, we concluded that 0.25 – 1 mg was the optimal range for SA-Mag beads for 0.5 mg of lysates. Given 1 mg of SA-Mag beads led to the greatest number of identified kinases without too much background binding and compromised kinase intensity, we selected this bead amount for subsequent experiments.

Prompted by the initial observation that linker length for biotinylated probes might affect target enrichment (Fig. 1B), we sought to assess the overall kinase capture efficiency of biotin-Cp19 probes with PEG2, 4, 8 or 12 linkers. Of the 142 kinases quantified across conditions, 0, 2, 1 and 3 were quantified uniquely by the PEG2, 4, 8 and 12 probe, respectively, while 94 were shared by them all (Fig. 2D, Table S3). In general, PEG8 and PEG12 linkers led to a greater overall intensity and a modestly increased number of quantified kinases than PEG2 and PEG4 (Fig. 2E, Fig. S1C), with PEG8 showing a slightly better overall kinase intensity. We therefore selected the Cp19-PEG8-biotin as our optimal probe.

We next compared the optimized automated AC workflow using SA-Mag (1 mg) and Cp19-PEG8-biotin (3.5 nmol) with the manual workflow using Cp19-Affi-Gel^™^ 10 (3.5 nmol effective Cp19 amount) for each sample using 0.5 mg cell lysate. As expected, the hands-on time needed for MS sample preparation was much shorter for the automated workflow (∼2 hours vs. ∼7 hours). We were additionally pleased to observe a similar distribution for protein intensity coefficient of variation (CV) between the two processes, with median CVs of 11.3% and 12.4% for the manual and automated workflows, respectively, reflecting robust protein quantification by both workflows (Fig. S2A). A reduced number of total kinases were quantified by the automated workflow (129) in comparison to the manual workflow (167), with a corresponding reduced number of uniquely detected kinases in the automated workflow (Fig. 2F, Table S4). Comparing the intensity for all proteins quantified revealed that the automated workflow yielded a lower overall intensity for kinases while comparable overall intensity and greater number for non-kinases than the manual workflow. As expected, streptavidin, trypsin and Lys-C were observed at high intensity in the SA-Mag samples, while only the latter two were detected in the Affi-Gel^™^ 10 samples (Fig. S2B). These results collectively suggest overall higher background identified by the automated workflow.

We reasoned that the inferior kinome coverage by the automated workflow could be attributed to the lower signal level for kinases and the higher background binding to SA-Mag. To overcome this limitation, we sought to exploit the data independent acquisition (DIA) method, which offers advantages over DDA schemes for characterizing complex protein digests with relatively short LC run times. In contrast to the sequential detection, selection, and analysis of individual ions during DDA, DIA parallelizes the fragmentation of all detectable ions within a wide m/z range regardless of intensity, thereby providing broader dynamic range of detected signals, improved reproducibility for identification, better sensitivity, and accuracy for quantification, and potentially enhanced proteome coverage^18, 19, 27^. As expected, DIA quantified 345 and 303 kinases with the manual and automated workflows using Cp19-based probes, which were >2-fold more kinases than DDA for both workflows (Fig. 2G, Table S5). Among all kinases quantified by DIA, 228 kinases were shared, corresponding to 66% and 75% of all kinases quantified by the manual and automated workflow, respectively. The overall good overlap in kinases between both approaches was in line with what was observed by DDA-MS (Fig. 2F). This number of overlapping kinases by DIA-MS matched a previous report for Cp19 that used 10 times the amount of cell lysate and a longer LC gradient with a kinase peptide-containing inclusion list^4^, giving us further confidence in the optimized workflow.

### Automated AC-MS workflow integrating DIA for kinome profiling

Encouraged by the boost in kinase coverage by DIA, we next opted to apply the optimized automated workflow to kinase inhibitor profiling by utilizing a combined panel of chemical probes and multiple cell lines for broad kinome coverage^4-6, 8, 28^. We selected the previously optimized multiprobe matrix KBγ comprising five kinase inhibitors^4^ and generated the corresponding biotin-tagged probes with PEG4 or PEG8 linkers (Fig. 3A). For improved kinome coverage^4^, we used a 1:1:1:1 mixture of lysates from K562, MV4-11, THP-1 and NCI-H1155 cells as the source of native kinases. Previous study showed that kinase coverage at the proteomic level could saturate at around four cell lines and that K562 and MV4-11 constitute a wide kinase repertoire^4^. Based on the amenability to large-scale cell culture and availability from the American Type Culture Collection (ATCC), we additionally included THP-1 and NCI-H1155, which, according to cell model passports^29^, afford 37 unique kinases in addition to the 237 detected in K562 and MV4-11 cell lines at the unenriched proteome level.

**Fig 3.**
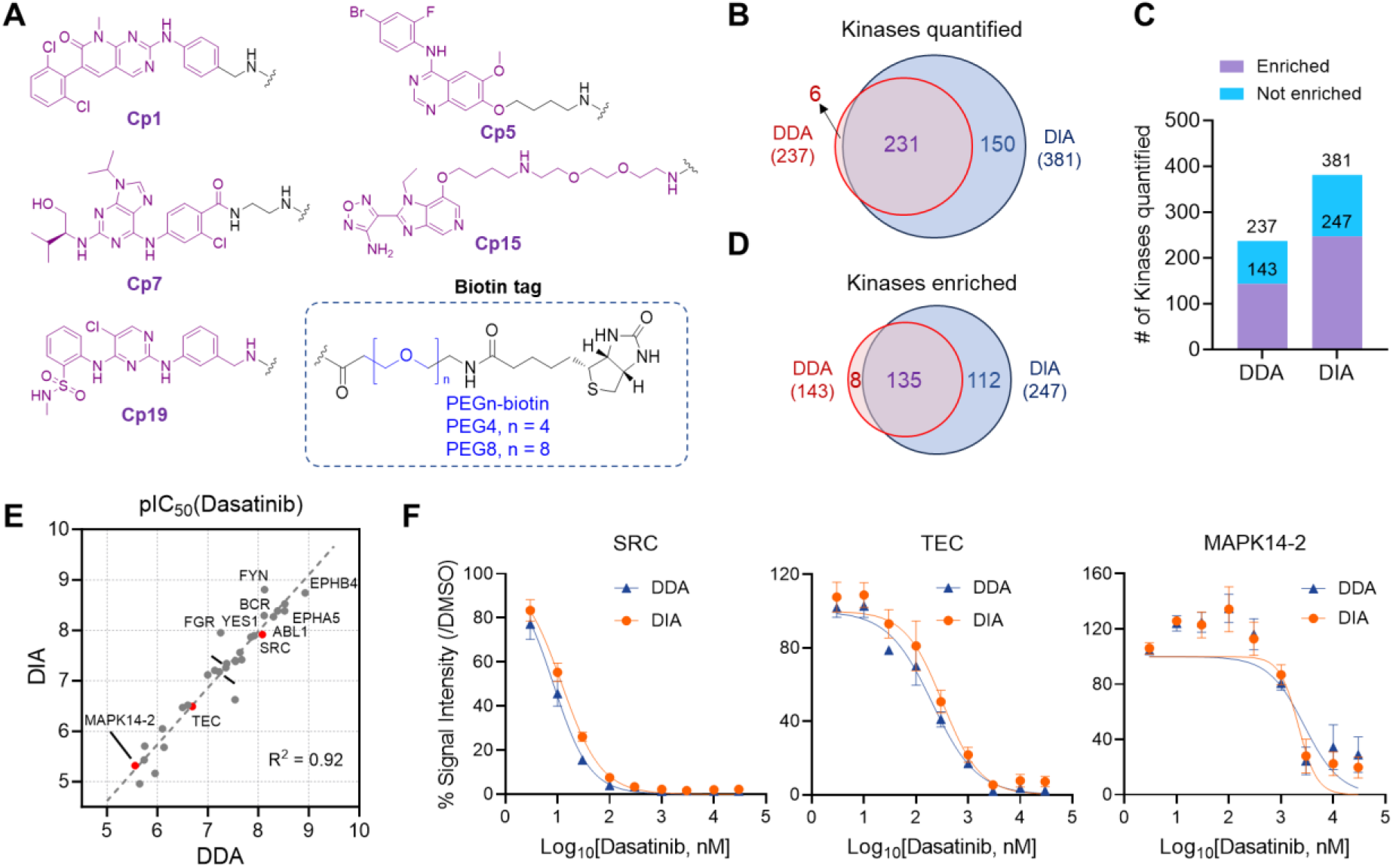
Automated AC-MS workflow using DIA for kinome profiling. (A) Structures for capturing compounds used to enrich kinases. (B, C, D) Venn diagram and bar plot showing number and overlap of kinases quantified and specifically enriched (≥2-fold change between with and without probe conditions, *P* < 0.05) in DDA- and DIA-MS analysis. All enrichments used equally mixed biotinylated affinity probes and mixed lysates of K562, MV4-11, THP-1 and NCI-H1155 cells. (E, F) Comparison of dasatinib pIC_50_ values calculated from DIA- and DDA-MS data across kinase targets (E) and normalized concentration-response data for representative targets (F). Red dots in (E) correspond to kinase targets shown in (F). Data in (F) are average values ± SEM. All experiments were performed with three independent replicates per group.

Comparing the intensity of quantified kinases using multiprobe matrices utilizing PEG4 and PEG8 linkers revealed greater kinase intensities with PEG8-containing probe set by both DDA- and DIA-MS (Fig. S3A). We thus focused on this set for the comparison between DDA and DIA. With probe enrichment, 237 and 381 kinases were quantified by DDA and DIA, respectively, with a 60.7% increase in kinase quantified by DIA compared with DDA (Table S6). Among all quantified kinases, 231 were common between both methods, 150 were uniquely quantified by DIA, while only 6 were uniquely quantified by DDA (Fig. 3B). Along with the boost in kinase coverage, we also noticed a 2.6× fold increase in total protein groups identified by DIA as compared with DDA (Fig. S3B). As expected, a higher percentage of low-intensity protein groups were found in those uniquely identified by DIA (Fig. S3C and D), which demonstrates the superiority of DIA in quantifying low-abundance peptides that DDA methods may miss^19, 27^. To further understand whether the increased kinase coverage was specific to those enriched by probes, we compared kinases quantified with versus without the biotinylated probes. 60.3% and 64.8% of all quantified kinases were specifically enriched (≥ 2x fold change, *P* ≤ 0.05) by the probes via DDA and DIA-MS analysis, respectively (Fig. 3C). Among the kinases enriched, 135 were common between both methods, with 112 uniquely enriched by DIA, while only 8 uniquely enriched by DDA (Fig. 3D).

Having established the ability of this probe set and automated AC-MS workflow to enrich ∼250 kinases, we next evaluated its ability to profile compound selectivity in a competition experiment. We selected the multi-kinase inhibitor dasatinib, which has been previously tested with analogous probe sets, by pre-incubating the mixture of cell extracts with nine concentrations from 0 to 30 µM before enrichment with the biotinylated probe set and SA-Mag beads. We identified 31 and 52 kinase targets (IC_50_ < 30 µM) for dasatinib by DDA- and DIA-MS, respectively. Intriguingly, all the DDA-identified dasatinib targets were identified by DIA, including 26 of previously identified targets via chemoproteomic kinome profiling, such as EPHA5, SRC, EPHB4, BCR and ABL1^5^ (Fig. S3E). The pIC_50_ values obtained for these targets determined from DDA and DIA-MS data additionally showed excellent correlation (R^2^ = 0.92, Fig. 3E). Furthermore, the concentration-response curves for three representative kinases, SRC, TEC and MAPK14, for which dasatinib had low, medium, and high IC_50_ values, tightly overlapped in normalized DDA and DIA data (Fig. 3F). Among the 21 targets uniquely identified using DIA-MS, 15 were shown to be targeted by dasatinib (Table S7) in previous kinome profiling^5^, further suggesting DIA-MS enabled identification of compound targets that could be overlooked by DDA-MS with short LC run time. Collectively, these results demonstrate the quantitative performance of the DIA workflow and its applicability to kinase inhibitor profiling.

### Kinome-wide selectivity of CDK9 inhibitors using modified affinity matrix

We next sought to employ our workflow to profile the selectivity of two related CDK9 inhibitors, BAY-1143572 (atuveciclib)^30^ and BAY-1251152 (VIP152, enitociclib)^31^, with the latter reported to exhibit better potency and improved therapeutic index than the former^31-33^. To explore potential improvements in kinome coverage and especially to capture CDK family members better, we first compared the kinases enriched by each of the KBγ probe individually. The Cp1, 15, and 19 probes each contributed 37, 12, and 32 uniquely enriched kinases, consistent with the previous reports that they are highly complementary^4^. On the other hand, the Cp5 did not enrich CDKs and the Cp5 and Cp7 probe enriched a relatively small number of unique and total number of kinases (Fig. 4A). We therefore decided to keep the Cp1, 15 and 19 probes and replace the Cp5 and 7 probes with a common kinase inhibitor scaffold Compound 8 (Cp8)^34^-derived probe and CDK inhibitor palbociclib (Palb)^35^-based probe (Fig. 4C). Cp8 and Palb probes enriched 191 and 122 kinases, respectively, among which 26 and 22 were unique compared to the ones covered by Cp1, 15 and 19 probes, including two additional CDKs, CDK8 and CDK19 (Fig. 4B, D, Table S8). The new five-probe combination was then tested in comparison to KBγ with identical quantity of each individual probe. The new matrix enriched 37 more kinases (Table S9) and led to enhanced enrichment for 71 kinases as compared to the KBγ set, including 6 CDK members, while only six kinases had reduced enrichment (Fig. 4E).

**Fig 4.**
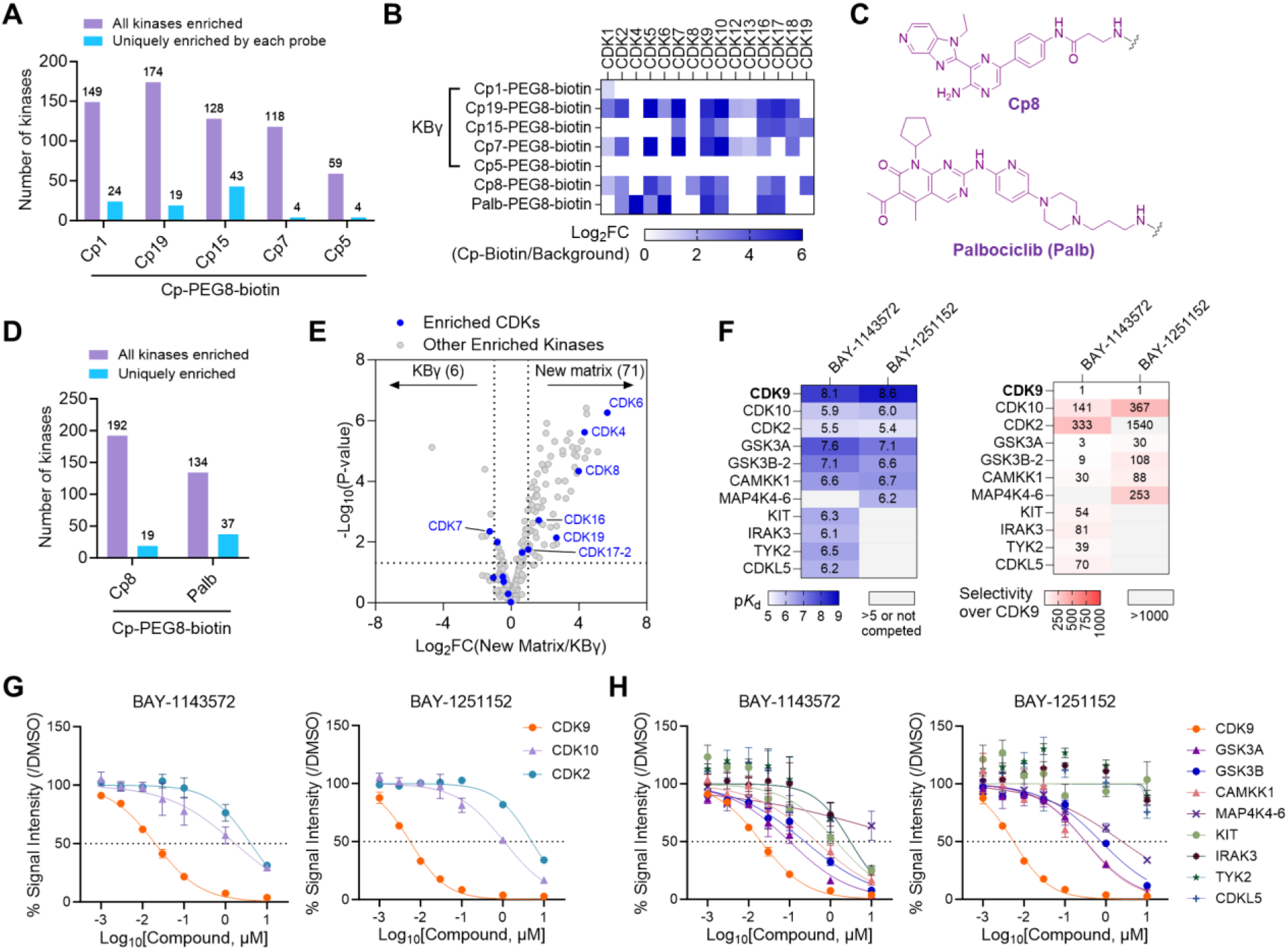
Kinome profiling of CDK9 inhibitors using a modified probe matrix. (A) Number of kinases significantly enriched by each KBγ probe individually. Uniquely enriched kinases are those only enriched by the indicated probe. Cutoff for enriched kinases: fold change (probe/background) ≥ 2, *P* < 0.05. (B) Heatmap showing enrichment of CDKs by indicated probes individually. Log_2_FC (probe/background) values are plotted. White color indicates CDK not quantified or not significantly enriched (*P* < 0.05 as significance cutoff). (C) Core chemical structures of 2 additional kinase capturing compounds, Cp8 and Palbociclib (Palb). (D) Number of kinases significantly enriched by Cp8 and Palb probes individually, as well as uniquely (not enriched by Cp1, 5 or 19 probes). (E) Comparison of MS intensities for kinases enriched by the new matrix containing Cp1, 5, 19, 8 and Palb probes vs. KBγ consisting of Cp1, 5, 19, 7 and 15. Kinases enriched by either matrix were plotted. (F) Heatmap showing p*K*_d_ values and fold selectivity for CDK9 for BAY-1143572 and BAY-1251152 against selected CDK kinases and other kinase targets. Kinases showing <1,000-fold CDK9 selectivity for either inhibitor were included. Fold selectivity was calculated by dividing *K*_d_ value for selected kinase by that for CDK9. (G, H) Concentration-response curves for CDK targets (G) and other selected kinase targets (H) of BAY-1143572 and BAY-1251152. Dotted line indicates 50% inhibition. Data in (G, H) are average values ± SEM. All experiments were performed with three independent replicates per group.

With the new matrix, we proceeded to profile the two CDK9 inhibitors to further demonstrate the general applicability and performance of the optimized AC-MS workflow. The inhibitors were tested at eight concentrations from 0 to 10 μM for a full concentration response. To allow conversion from IC_50_ to *K*_d_ for the interaction of each kinase with the inhibitors, we included a second enrichment step to account for bead-matrix induced depletion of kinases from equilibrium^4-6, 28^. As expected, both BAY-1143572 and BAY-1251152 are highly potent and selective for CDK9 over the other 13 CDKs profiled in our study. Consistent with previous reports, BAY-1251152 exhibited better potency towards CDK9 and greater selectivity against the structurally related CDK2 than BAY-1143572 (Fig. 4F, Table S10, ratio of *K*_d_ values CDK2/CDK9 for BAY-1143572 vs. BAY-1251152: 333 vs. 1,547). Interestingly, our results indicated that both compounds also weakly engaged another CDK family member, CDK10, and exhibited a slightly greater potency for CDK10 over the known off-target CDK2 (Fig. 4F and 4G). Leveraging that the workflow enriches kinases along with their interactors^36^, we observed that both compounds also displaced the enrichment for the corresponding cyclin partners of CDK9, 2 and 10 – cyclin T1/T2 (CCNT1/T2), cyclin M (CCNQ) and cyclin E2 (CCNE2) (Fig. S4A)– correlating with their potency against each of the CDKs (Fig. S4B). These results nicely cross-validated the compounds’ effects on the CDKs.

Outside the CDK family, seven and four kinases, respectively, were targeted by BAY-1143572 and BAY-1251152 with a *K*_d_ ratio less than 1,000 (GSK3A, GSK3B and CAMKK1 for both compounds, KIT, IRAK3, TYK2 and CDKL5 for BAY-1143572 only, and MAP4K4 for BAY-1251152 only). Based the *K*_d_ ratio values, BAY-1251152 was more selective for CDK9 than BAY-1143572 over all potential kinase off-targets identified except for MAP4K4 (Fig. 4F and H). The three overlapping kinases, GSK3A, GSK3B and CAMKK1, appeared to be the top three off-targets for both compounds. Among them, GSK3A and GSK3B have been reported to be targeted and inhibited by both compounds^31, 33^, while CAMKK1 (*K*_d_ for BAY-1143572 and BAY-1251152: 0.27 and 0.22 µM) was shown to be inhibited by BAY-1251152 in DiscoverX Kinome Scan data (87% at 1 µM)^33^. Additionally, MAP4K4 as a weak off-target for BAY-1251152 (*K*_d_ value 0.62 μM) was also consistent with previously reported DiscoverX data (56% inhibition at 1 µM^33^). KIT, IRAK3, TYK2 and CDKL5 (*K*_d_ values of 0.48, 0.72, 0.34 and 0.62 μM, respectively), however, have not been previously reported as potential off-targets for BAY-1143572 possibly due to the lack of comprehensive profiling. Collectively, the data presented here provides a view of the selectivity of two highly related CDK9 inhibitors against native kinome that is complementary to in vitro measurements^30, 31, 33^ and the cellular NanoBRET assay^33, 37^ and demonstrates the utility of the streamlined AC-MS workflow to comprehensively characterize kinase inhibitor selectivity in the native proteome.

## Discussion

In this study we describe the development, optimization, and application of a streamlined and automated AC-MS workflow, which enables chemoproteomic kinome profiling in a high-throughput, robust and time-effective manner. Using biotin-conjugated probes in combination with magnetic beads, we removed the requirement for the laborious centrifugation- or filter plate-based steps to separate unbound proteome and exchange sample buffers from beads, as well as enable enrichment, wash, reduction, alkylation and digestion steps in a one-pot, 96-well plate format. This workflow was automated using a pipetting robot, significantly reducing hands-on time, and minimizing potential human errors inherent in manual steps. The throughput of MS analysis was optimized by employing an EvoSep LC system alongside label-free DIA-MS methods to achieve instrument runtimes of 44 min per sample. The label-free approach offers several advantages, including enabling comparison among hundreds of samples and eliminating costs associated with isotope labeling reagents.

We observed that magnetic beads we use nonspecifically bind more proteins than the Affi-Gel^™^ 10-based agarose beads, to the detriment of kinase coverage in a DDA-MS method. By integrating DIA-MS, which bypasses the stochastic nature of DDA-MS by sampling and fragmenting all precursors within predefined *m*/*z* windows, we substantially improve the kinases quantified and enriched by >60% in our updated workflow. The resulting number of quantified kinases was superior to those previously reported using Sepharose immobilized affinity matrix with 20×, 5× to later miniaturized amount of cell lysates and longer LC gradient^4, 5, 8^. The observed nonspecific binding prompted us to evaluate kinases pulled down independent of biotinylated probes: >30% of quantified kinases showed <2-fold enrichment in the presence of probes by both DDA- and DIA-MS, excluding them from the pool of kinases that can be assayed through this chemoproteomic approach. Therefore, evaluation of background binding is highly recommended for future studies using biotinylated probes and magnetic beads to avoid drawing conclusions for proteins that are non-specifically pulled down.

Automation is being continuously employed to increase throughput and robustness for proteomic^38-40^ and chemoproteomic^6, 14, 41^ applications. Herein we demonstrated the use of biotinylated small molecule probes for automation by pre-incubating the probes with magnetic beads. We envision the established workflow could serve as a template for affinity capture of protein families beyond kinases using broad-spectrum chemical probes, such as those that bind histone deacetylases^42^, or bromodomain proteins^10^. This approach can also be used to identify protein targets specific to a small molecule of interest using more tailored chemical probes. Our automated workflow was set up using INTEGRA Assist Plus to process up to an entire 96-well plate of samples at once but should be straightforward to implement for users with access to other systems utilizing magnetic separations, such as the KingFisher system (Thermo Fisher Scientific Scientific). The magnetic-based protocol could alternatively be performed manually with a multichannel pipet and magnetic rack, which could be a viable option to skip the cumbersome centrifugation-assisted washing steps.

We used our optimized workflow to profile two related clinical stage CDK9 inhibitors, BAY-1143572 and BAY-1251152. Gratifyingly, our study reproduced the previously reported differential potency and selectivity for CDK9 between the two compounds and their off-target binding to CDK2, GSK3A, GSK3B, CAMKK1 or MAP4K4 among the ∼250 kinases enriched in our study^31, 33, 37^. Our study also revealed a previously unknown weak off-target for BAY-1251152, CDK10 (*K*_d_ 0.90 μM), which remains one of the most elusive members of the CDK family^43^. CDK10 is not in the DiscoverX 468-kinase Kinome Scan panel and therefore was not tested in the previous kinome selectivity profiling for BAY-1251152^33^. In the study where CDK inhibitor selectivity was profiled in live cells using energy-transfer probes^37^, CDK10 was assayed in complex with cyclin L2 rather than cyclin M, which we identified as the cyclin partner blocked by BAY-1251152. Considering engagement of CDK for some compounds could shift dramatically depending on the cyclin partner^37^, it is possible that BAY-1251152 targets the CDK10/cyclin M complex more potently than CDK10/cyclin L2. This observation highlights that the AC-MS approach enables quantification of target occupancy of native kinases without characterization and functional annotation. While our data suggest that BAY-1251152 targets several other kinases, it still has >30∼1,000-fold selectivity for CDK9. To what extent these off-targets are pharmacologically relevant remains to be investigated.

We finally conclude by emphasizing the potential of the magnetic bead-based automated high-throughput AC-MS platform. The advancements in ultra-fast and sensitive MS instrumentation^19, 44, 45^ points to the possibility to perform affinity capture with further miniaturization and to reduce the LC gradient to 11 min per sample (100 samples per day) or even shorter run times. The optimized sample preparation protocol outlined here can be readily tailored for smaller proteome quantities and adapted to a 384-well plate format, which we envision would facilitate the analysis of precious sample types such as primary cells or enable large-scale screening for profiling compound selectivity against kinases and other protein classes.

## Supporting information

Supplemental figures and chemical synthesis

DIAWindow_tMSn_MassLists

## AbbVie Disclosure Statement

H. J., G.K.P., S.S., V.L.M., A.B., H.T., J. E.K., R.M., J.D.W. are employees of AbbVie. P.L.R and R. M. were employees of AbbVie at the time of the study. The design, study conduct, and financial support for this research were provided by AbbVie. AbbVie participated in the interpretation of data, review, and approval of the publication.

