## Supplemental figures and chemical synthesis for "An Automated High-throughput Affinity Capture-Mass Spectrometry Platform with Data- Independent Acquisition"

[1] Discovery Research, AbbVie, Inc., 1 North Waukegan Rd., North Chicago, Illinois 60064, United States.
[#]Contributed equally.

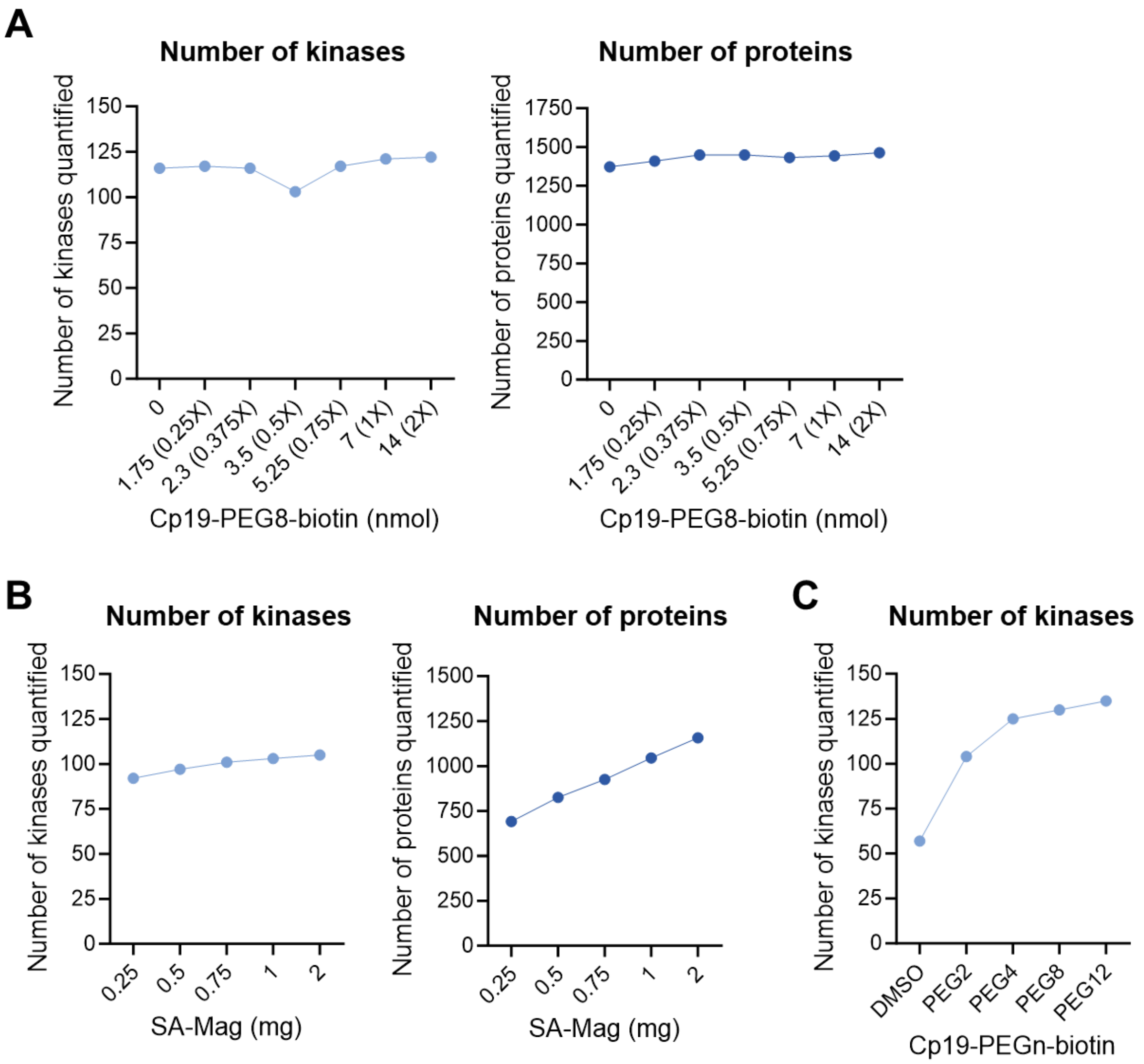


**Figure. S1** Impact of the amount of (A) Cp19-biotin, (B) SA-Mag beads and (C) PEG linker length on the number of kinases and total proteins quantified. (A, B) Number of kinases and proteins quantified by using (A) a fixed amount of SA-Mag beads (2 mg) and varying biotin-PEG8-Cp19 loading or (B) an indicated amount of SA-Mag at a fixed a probe/bead ratio of 3.5 nmol/mg. (C) Number of kinases quantified by using Cp19-biotin probes with PEG2, 4, 8 and 12 linkers.

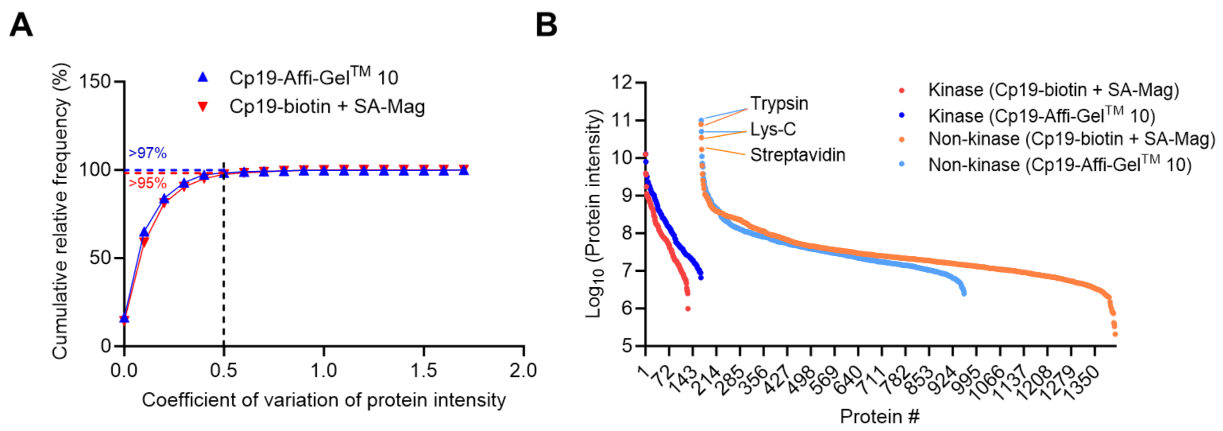


**Figure. S2** Comparison between the automated workflow using Cp19-PEG8-biotin and the manual workflow using Cp19-Affi-Gel$^{TM}$ 10. (A) Cumulative relative frequency of protein intensity coefficient of variation (CV) by the automated workflow using Cp19-PEG8-biotin in combination with SA-Mag beads and the manual workflow using Cp19-Affi-Gel$^{TM}$ 10. >95% and >97% indicate the relative frequency of proteins with <0.5 CV for the automated and manual workflow, respectively. (B) Protein rank plot showing intensity and number of kinases and non-kinase proteins quantified by the automated and manual workflows. Plotted are median Log10(LFQ intensity) values for each protein across three replicates.

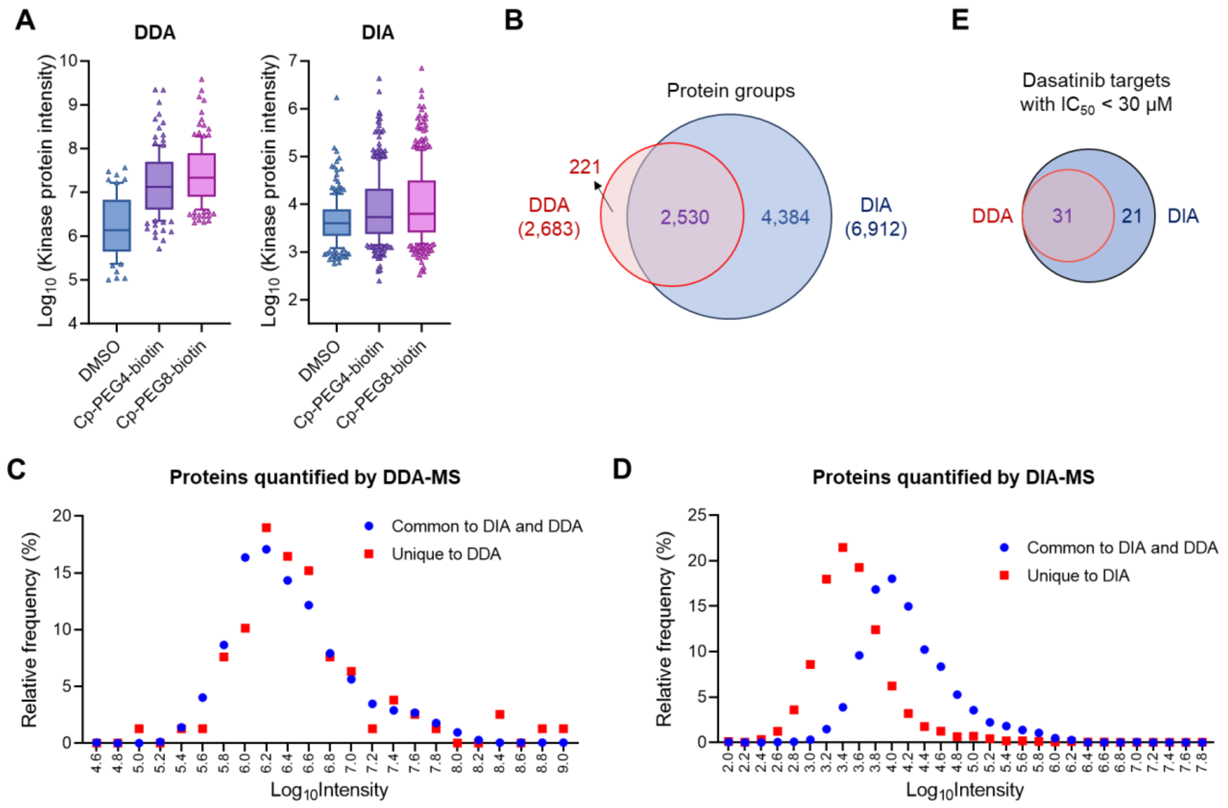


**Figure. S3** Comparison between DDA-MS and DIA-MS in automated kinome profiling. (A) MS intensities for kinases enriched using mixed biotinylated probes with PEG4 or PEG8 linkers and quantified by DDA- and DIA-MS. Cp1, 5, 7, 15, and 19-biotin probes with the same linker lengths were mixed 1:1:1:1:1 to a total amount of 3.5 nmol and pre-incubated with 1 mg of SA-Mag per sample. (B) Venn diagram indicating the number of protein groups quantified by DDA- and DIA-MS with the PEG8 linker-containing probe mixture. (C, D) Intensity distribution for protein groups quantified by (C) DDA-MS and (D) DIA-MS. Protein groups common to DIA and DDA, and unique to DDA (C) or DIA (D) were plotted. (E) Venn diagram showing the overlap of dasatinib targets identified by DDA- and DIA-MS. Kinases with significantly inhibited enrichment at 30 μM ($P < 0.05$) and $IC_{50} < 30$ μM were counted.

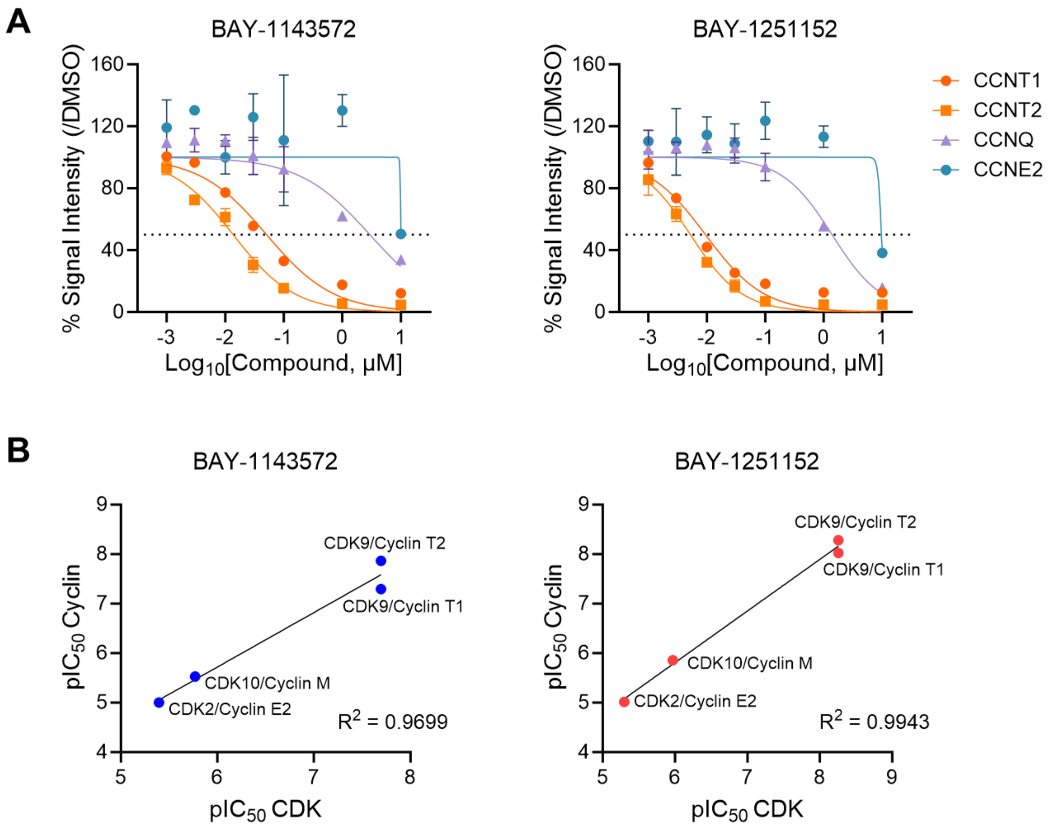


**Figure. S4** Displacement of cyclin proteins by the CDK9 inhibitors, BAY-1143572 and BAY-1251152. (A) Concentration-response curves for cyclin T1 (CCNT1), cyclin T2 (CCNT2), cyclin M (CCNQ) and cyclin E2 (CCNE2) of BAY-1143572 and BAY-1251152. Dotted line indicates 50% inhibition. (B) Correlation of $pIC_{50}$ values for BAY-1143572 or BAY-1251152 against CDKs and the corresponding cyclin partners. $R^2$ values for liner correlation are shown.

**Chemical Synthesis**

**Reagents:** Unless otherwise specified all the reagents and solvents used for the synthesis of compounds were purchased from Millipore-Sigma, St. Louis MO. 2,5-Dioxopyrrolidin-1-yl 17-oxo-21-((3aS,4S,6aR)-2-oxohexahydro-1H-thieno[3,4-d]imidazol-4-yl)-4,7,10,13-tetraoxa-16-azahenicosan-1-oate (biotin-PEG4-NHS), 2,2-dimethyl-4-oxo-3,8,11-trioxa-5-azatridecan-13-oic acid, 2,5-dioxopyrrolidin-1-yl 5-((3aS,4S,6aR)-2-oxohexahydro-1H-thieno[3,4-d]imidazol-4-yl)pentanoate (biotin-NHS), *tert*-butyl (3-oxopropyl) and 7-Azabenzotriazol-1-yloxy)tripyrrolidinophosphonium hexafluorophosphate (PyAOP) were obtained from Combi-Blocks, San Diego, CA. 2,5-Dioxopyrrolidin-1-yl 29-oxo-33-((3aS,4S,6aR)-2-oxohexahydro-1H-thieno[3,4-d]imidazol-4-yl)-4,7,10,13,16,19,22,25-octaoxa-28-azatritriacontan-1-oate (biotin-PEG8-NHS), N-(26-amino-3,6,9,12,15,18,21,24-octaoxahexacosyl)-5-((3aS,4S,6aR)-2-oxohexahydro-1H-thieno[3,4-d]imidazol-4-yl)pentanamide (biotin-PEG8-$NH_2$), 2,5-dioxopyrrolidin-1-yl 41-oxo-45-((3aS,4S,6aR)-2-oxohexahydro-1H-thieno[3,4-d]imidazol-4-yl)-4,7,10,13,16,19,22,25,28,31,34,37-dodecaoxa-40-azapentatetracontanoate (biotin-PEG12-NHS), 2,5-dioxopyrrolidin-1-yl 23-oxo-27-((3aS,4S,6aR)-2-oxohexahydro-1H-thieno[3,4-d]imidazol-4-yl)-4,7,10,13,16,19-hexaoxa-22-azaheptacosan-1-oate (biotin-PEG6-NHS) and 2,5-dioxopyrrolidin-1-yl 3-(2-(2-(5-((3aS,4S,6aR)-2-oxohexahydro-1H-thieno[3,4-d]imidazol-4-yl)pentanamido)ethoxy)ethoxy)propanoate (biotin-PEG2-NHS) were obtained from Broadpharm, San Diego, CA. Purvalinol B was obtained from Synnovator, Inc., Cary, NC. Affi-Gel™10-NHS affinity resin was obtained from Bio-Rad, Hercules, CA.

**Abbreviations:**

| | |
|---|---|
| LCMS | liquid chromatography-mass spectrometry |
| HATU | 1-[Bis(dimethylamino)methylene]-1H-1,2,3-triazolo[4,5-b]pyridinium 3-oxid hexafluorophosphate |
| BME | 2-mercaptoethanol |
| DBU | DBU (1,8-diazabicyclo[5.4.0]undec-7-ene) |
| HPLC | high-performance liquid chromatography |
| RP-HPLC | reverse-phase high-performance liquid chromatography |
| APCI-MS | atmospheric pressure chemical ionization mass spectrometry |
| ESI-MS | electrospray ionization mass spectrometry |
| TFA | trifluoroacetic acid |
| $R_t$ | Retention time |
| AA | ammonium acetate |
| FA | formic acid |
| NMR | nuclear magnetic resonance |
| DMSO | dimethylsulfoxide |
| COSY | correlation spectroscopy |
| HSQC | heteronuclear single quantum coherence spectroscopy |
| HMBC | heteronuclear multiple-bond correlation spectroscopy |
| ROESY | rotating frame nuclear Overhauser effect spectroscopy |
| DMF | dimethylformamide |
| DIEA | diisopropylethylamine |
| MeOH | methanol |
| THF | tetrahydrofuran |
| DCM | dichloromethane |
| psi | per square inch |
| PyAOP | 7-Azabenzotriazol-1-yloxy)tripyrrolidinophosphonium hexafluorophosphate |

| | |
|---|---|
| biotin-NHS | 2,5-dioxopyrrolidin-1-yl 5-((3aS,4S,6aR)-2-oxohexahydro-1H-thieno[3,4-d]imidazol-4-yl)pentanoate |
| biotin-PEG2-NHS | 2,5-dioxopyrrolidin-1-yl 3-(2-(2-(5-((3aS,4S,6aR)-2-oxohexahydro-1H-thieno[3,4-d]imidazol-4-yl)pentanamido)ethoxy)ethoxy)propanoate |
| biotin-PEG4-NHS | 2,5-Dioxopyrrolidin-1-yl 17-oxo-21-((3aS,4S,6aR)-2-oxohexahydro-1H-thieno[3,4-d]imidazol-4-yl)-4,7,10,13-tetraoxa-16-azahenicosan-1-oate |
| biotin-PEG6-NHS | 2,5-dioxopyrrolidin-1-yl 23-oxo-27-((3aS,4S,6aR)-2-oxohexahydro-1H-thieno[3,4-d]imidazol-4-yl)-4,7,10,13,16,19-hexaoxa-22-azaheptacosan-1-oate |
| biotin-PEG8-NHS | 2,5-Dioxopyrrolidin-1-yl 29-oxo-33-((3aS,4S,6aR)-2-oxohexahydro-1H-thieno[3,4-d]imidazol-4-yl)-4,7,10,13,16,19,22,25-octaoxa-28-azatritriacontan-1-oate |
| biotin-PEG12-NHS | 2,5-dioxopyrrolidin-1-yl 41-oxo-45-((3aS,4S,6aR)-2-oxohexahydro-1H-thieno[3,4-d]imidazol-4-yl)-4,7,10,13,16,19,22,25,28,31,34,37-dodecaoxa-40-azapentatetracontanoate |
| biotin-PEG8-$NH_2$ | N-(26-amino-3,6,9,12,15,18,21,24-octaoxahexacosyl)-5-((3aS,4S,6aR)-2-oxohexahydro-1H-thieno[3,4-d]imidazol-4-yl)pentanamide |

**Synthetic Methods**

**Analytical LCMS with FA method 1 and AA method 1:** Analytical LCMS was performed on a Thermo MSQ-Plus mass spectrometer and Agilent 1100/1200 HPLC system running Xcalibur 2.0.7, Open-Access 1.4, and custom login software. The HPLC system comprised an Agilent Binary pump, degasser, column compartment, autosampler and diode-array detector, with a Polymer Labs ELS-2100 evaporative light-scattering detector. The column used was a Waters Cortecs C18+, 2.7 µm 100Å (2.1mm × 30mm), at a temperature of 40°C. Elution times are reported as $R_t$.

**FA method 1:** A gradient of 5-100% acetonitrile (A) and 0.1% formic acid water:acetonitrile (98:2) (B) was used, at a flow rate of 1.5 mL/min (0-0.05 min 0% A, 0.05-2.8 min 0-100% A, 2.8-3.0 min 100-0% A, 160-1500 amu positive/negative ESI-MS ionization. Elution times are reported as $R_t$.

**AA method 1:** A gradient of 5-100% acetonitrile (A) and 10 mM ammonium acetate water:acetonitrile (98:2) (B) was used, at a flow rate of 1.5 mL/min (0-0.05 min 0% A, 0.05-2.8 min 0-100% A, 2.8-3.0 min 100-0% A, 160-1500 amu positive/negative ESI-MS ionization. Elution times are reported as $R_t$.

**TFA method 1:** Analytical LCMS was performed on a Thermo MSQ-Plus mass spectrometer and Agilent 1100/1200 HPLC system running Xcalibur 2.0.7, Open-Access 1.4, and custom login software. The mass spectrometer was operated under positive APCI or ESI ionization conditions dependent on the system used, as noted in the text. The HPLC system comprised an Agilent Binary pump, degasser, column compartment, autosampler and diode-array detector, with a Polymer Labs ELS-2100 evaporative light-scattering detector. The column used was a Phenomenex Kinetex Cp8, 2.6 µm 100 Å (2.1mm × 30mm), at a temperature of 65°C. A gradient of 5-100% acetonitrile (A) and 0.1% trifluoroacetic acid in water (B) was used, at a flow rate of 1.5 mL/min (0-0.05 min 5% A, 0.05-1.2 min 5-100% A, 1.2-1.4 min 100% A, 1.4-1.5 min 100-5% A. 0.25 min post-run delay). Elution times are reported as $R_t$.

**Preparative HPLC TFA method 1:** RP-HPLC was performed on a Gilson HPLC system equipped with liquid handler (Gilson-215) and UV/Vis detector using a Waters Deltapak C18 column (5 µm, 100 Å, 200 x 21.2 mm) eluted with A (0.1% TFA-water):B (acetonitrile) [0-5 min: 5% A; 5-35 min linear gradient to 100% B, 3.16%/min linear gradient] with a 20 mL/min flowrate.

**Preparative HPLC TFA method 2:** RP-HPLC was performed on a Gilson HPLC system equipped with liquid handler (Gilson-215) and UV/Vis detector using a Phenomenex Gemini NX-C18 column (5 µm, 110 Å, 250 x 25 mm) eluted with A (0.1% TFA-water):B (acetonitrile) [0-5 min: 2% A; 5-35 min linear gradient to 100% B, 3.16%/min linear gradient] with a 20 mL/min flowrate.

**NMR** Unless otherwise specified all spectra were collected in DMSO-$d_6$ at room temperature (27°C) on a Bruker Avance III HD spectrometer equipped with a TCI cryoprobe. $^1$H spectra were acquired with SW= 16 ppm, 4.1 s acquisition time & 24 scans.

**2-((4-(Aminomethyl)phenyl)amino)-6-(2,6-dichlorophenyl)-8-methylpyrido[2,3-d]pyrimidin-7(8H)-one (Cp1)** was synthesized by WuXi Apptec, Shanghai, China as described (1)(2).

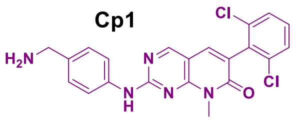


LCMS FA method 1: $R_t$ = 0.994 min, ESI-MS *m/z* 426.0 $(M+H)^+$, 423.8 $(M-H)^-$; LCMS AA method 1: $R_t$ = 1.302 min, ESI-MS *m/z* 426.0 $(M+H)^+$, 424.0 $(M-H)^-$; $^1H$ NMR (400 MHz, DMSO-$d_6$) $\delta_H$ 10.28 (s, 1H), 8.85 (s, 1H), 7.91 (s, 1H), 7.82 (d, *J* = 8.4 Hz, 2H), 7.59 (d, *J* = 8.0 Hz, 2H), 7.47 (t, *J* = 8.2 Hz, 1H), 7.39(d, *J* = 8.4 Hz, 2H), 6.06 (br, 2H), 3.87 (s, 2H), 3.67 (s, 3H).


(1) Klutchko, S.R. et al, *J. Med. Chem.,* **1998**, 41(17), 3276-3292.
(2) Baldwin, I.R. et al, PCT Int. Appl., **2008**, WO 2008110508 A1 page 49.


**N-(4-((6-(2,6-dichlorophenyl)-8-methyl-7-oxo-7,8-dihydropyrido[2,3-d]pyrimidin-2-yl)amino)benzyl)-1-(5-((3aS,4S,6aR)-2-oxohexahydro-1H-thieno[3,4-d]imidazol-4-yl)pentanamido)-3,6,9,12-tetraoxapentadecan-15-amide 2,2,2-trifluoroacetate (Cp1-PEG4-biotin):**

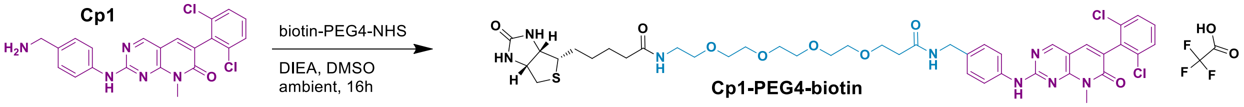


To 2-((4-(Aminomethyl)phenyl)amino)-6-(2,6-dichlorophenyl)-8-methylpyrido[2,3-d]pyrimidin-7(8H)-one (Cp1, 54.3 mg, 0.127 mmol) and biotin-PEG4-NHS (50 mg, 0.085 mmol) dissolved in 1 mL anhydrous DMSO was added DIEA (44 mg, 0.34 mmol). The reaction was shaken at ambient overnight. The reaction was diluted to 3 mL with 90% DMSO/water (v/v) and purified by preparative HPLC TFA method 1. Fractions containing the desired peak were combined and lyophilized to give the desired product **Cp1-PEG4-biotin** (67.0 mg, 77.9 %) as a colorless waxy solid: LCMS FA method 1: $R_t$ = 1.430 min, ESI-MS *m/z* 450.2 $(M+2H)^{2+}$, 901.2 $(M+H)^+$, 899.2 $(M-H)^-$; LCMS AA method 1: $R_t$ = 1.451 min, ESI-MS *m/z* 450.2 $(M+2H)^{2+}$, 901.2 $(M+H)^+$, 899.0 $(M-H)^-$; $^1H$ NMR (600 MHz, DMSO-$d_6$) $\delta_H$ 10.19 (s, 1H), 8.83 (s, 1H), 8.33 (t, *J* = 5.9 Hz, 1H), 7.88 (s, 1H), 7.82 (t, *J* = 5.7 Hz, 1H), 7.77 (d, *J* = 8.2 Hz, 2H), 7.57 (d, *J* = 8.1 Hz, 2H), 7.45 (dd, *J* = 8.6, 7.6 Hz, 1H), 7.24 (d, *J* = 8.4 Hz, 2H), 6.42 (s, 2H), 4.30 (ddd, *J* = 7.8, 5.2, 1.0 Hz, 1H), 4.25 (d, *J* = 5.9 Hz, 2H), 4.12 (dd, *J* = 7.8, 4.4 Hz, 1H), 3.65 (d, *J* = 6.9 Hz, 5H), 3.54 – 3.44 (m, 12H), 3.17 (q, *J* = 5.8 Hz, 2H), 3.08 (ddd, *J* = 8.6, 6.1, 4.4 Hz, 1H), 2.80 (dd, *J* = 12.5, 5.1 Hz, 2H), 2.58 (d, *J* = 12.4 Hz, 1H), 2.39 (t, *J* = 6.3 Hz, 2H), 2.05 (t, *J* = 7.4 Hz, 2H), 1.53 – 1.40 (m, 2H), 1.34 – 1.24 (m, 2H); $^{13}C$ NMR (151 MHz, DMSO-$d_6$) $\delta_C$ 172.2, 170.12, 162.69, 160.54, 159.22, 158.92, 155.21, 137.96, 136.86, 134.94 (3 C), 133.62, 130.63, 128.02 (2 C), 127.5 (2 C), 124.1, 119.66 (2 C), 105.64, 69.68 (4 C), 69.52, 69.5, 69.02, 66.79, 61.07, 59.14, 55.27, 41.6, 39.76, 38.39, 36.12, 35.0, 28.1, 27.97, 27.85, 25.18.

**N-(4-((6-(2,6-dichlorophenyl)-8-methyl-7-oxo-7,8-dihydropyrido[2,3-d]pyrimidin-2-yl)amino)benzyl)-1-(5-((3aS,4S,6aR)-2-oxohexahydro-1H-thieno[3,4-d]imidazol-4-yl)pentanamido)-3,6,9,12,15,18,21,24-octaoxaheptacosan-27-amide 2,2,2-trifluoroacetate (Cp1-PEG8-biotin):**

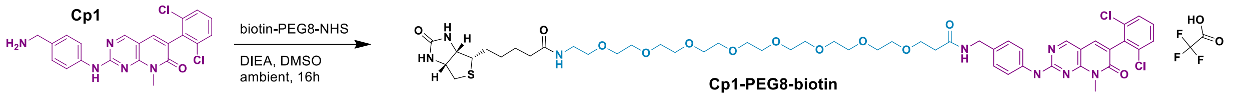

To 2-((4-(Aminomethyl)phenyl)amino)-6-(2,6-dichlorophenyl)-8-methylpyrido[2,3-d]pyrimidin-7(8H)-one (Cp1, 21.8 mg, 0.051 mmol) and biotin-PEG8-NHS (39.2 mg, 0.051 mmol) dissolved in 1 mL anhydrous DMSO was added DIEA (30 mg, 0.230 mmol). The reaction was shaken at ambient overnight. The reaction was diluted to 3 mL with 90% DMSO/water (v/v) and purified by preparative HPLC TFA method 2. Fractions containing the desired peak were combined and lyophilized to give the desired product **Cp1-PEG8-biotin** (39.2 mg, 64.3 %) as a colorless waxy solid: LCMS FA method 1: $R_t$ = 1.456 min, ESI-MS *m/z* 538.2 $(M+2H)^{2+}$, 1077.0 $(M+H)^+$, 1073.2 $(M-H)^-$; LCMS AA method 1: $R_t$ = 1.472 min, ESI-MS *m/z* 539.2 $(M+2H)^{2+}$, 1075.2 $(M+H)^+$, 1073.2 $(M-H)^-$; $^1$H NMR (600 MHz, DMSO-$d_6$) δ 10.21 (s, 1H), 8.83 (s, 1H), 8.31 (t, *J* = 5.9 Hz, 1H), 7.89 (s, 1H), 7.81 (s, 0H), 7.78 (d, *J* = 8.3 Hz, 2H), 7.60 – 7.55 (m, 2H), 7.45 (dd, *J* = 8.6, 7.6 Hz, 1H), 7.27 – 7.22 (m, 2H), 6.42 (s, 2H), 4.30 (m, 1H), 4.26 (d, *J* = 5.9 Hz, 2H), 4.12 (dd, *J* = 7.7, 4.4 Hz, 1H), 3.65 (d, *J* = 8.1 Hz, 5H), 3.54 – 3.45 (m, 28H), 3.39 (t, *J* = 5.9 Hz, 2H), 3.21 – 3.15 (m, 3H), 3.08 (m, 1H), 2.81 (dd, *J* = 12.4, 5.1 Hz, 1H), 2.58 (d, *J* = 12.4 Hz, 1H), 2.39 (t, *J* = 6.4 Hz, 2H), 2.06 (t, *J* = 7.4 Hz, 2H), 1.61 (m, 1H), 1.55 – 1.41 (m, 3H), 1.35 – 1.25 (m, 2H); $^{13}$C NMR (151 MHz, DMSO-$d_6$) $\delta_C$ 172.1, 170.02, 162.64, 160.55, 159.19, 158.87, 155.19, 138.04, 136.82, 134.89 (2 C), 134.29, 133.71, 130.53, 128.05 (2 C), 127.53 (2 C), 124.01, 119.65 (2 C), 105.6, 69.62 (12 C), 69.50, 56.49, 69.13, 66.85, 61.06, 59.13, 55.37, 41.51, 39.77, 38.35, 36.12, 35.05, 38.10, 27.91, 27.89, 25.14.

**7-(4-Aminobutoxy)-N-(4-bromo-2-fluorophenyl)-6-methoxyquinazolin-4-amine bis 2,2,2-trifluoroacetate (Cp5)** was synthesized as described (3) except the final product was dissolved in 90% DMSO/water (v/v) and purified by preparative HPLC TFA method 1. Fractions containing the desired peak were combined and lyophilized to give the desired product **Cp5** as a colorless powder.

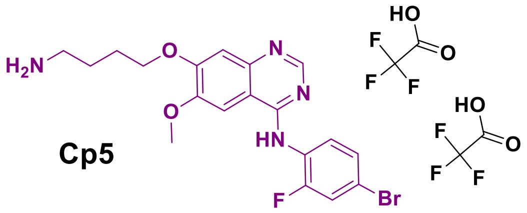

LCMS FA method 1: $R_t$ = 0.71 min, ESI-MS *m/z* 219.0 $(M+2H)^{2+}$, 435.0 $(M+H)^+$, 433.0 $(M-H)^-$; LCMS AA method 1: $R_t$ = 1.281 min (broad peak), ESI-MS *m/z* 437.0 $(M+H)^+$, 434.8 $(M-H)^-$; $^1$H NMR (600 MHz, PYRIDINE) $\delta_H$ 7.90 (s, 1H), 7.85 (t, *J* = 8.4 Hz, 1H), 7.58 – 7.53 (m, 3H), 7.48 (dd, *J* = 9.9, 2.2 Hz, 1H), 7.36 (ddd, *J* = 8.6, 2.3, 1.0 Hz, 1H), 4.11 (t, *J* = 6.0 Hz, 2H), 3.62 (s, 3H), 3.38 (dd, *J* = 8.2, 6.8 Hz, 2H), 2.20 – 2.12 (m, 2H), 1.98 (dt, *J* = 8.2, 6.1 Hz, 2H).

(3) Drewes, G. et al, **2006**, WO 2006134056 A1 page 100.

**N-(4-((4-((4-bromo-2-fluorophenyl)amino)-6-methoxyquinazolin-7-yl)oxy)butyl)-1-(5-((3aS,4S,6aR)-2-oxohexahydro-1H-thieno[3,4-d]imidazol-4-yl)pentanamido)-3,6,9,12-tetraoxapentadecan-15-amide 2,2,2-trifluoroacetate (Cp5-PEG4-biotin):**

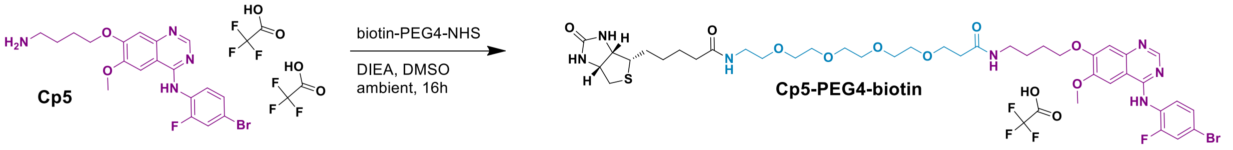


To 7-(4-Aminobutoxy)-N-(4-bromo-2-fluorophenyl)-6-methoxyquinazolin-4-amine bis-2,2,2-trifluoroacetate (**Cp5**, 56.3 mg, 0.085 mmol) biotin-PEG4-NHS (50 mg, 0.085 mmol) dissolved in 1 mL anhydrous DMSO was added DIEA (44 mg, 0.34 mmol). The reaction was shaken at ambient overnight. The reaction was diluted to 3 mL with 90% DMSO/water (v/v) and purified by preparative HPLC TFA method 1. Fractions containing the desired peak were combined and lyophilized to give the desired product **Cp5-PEG4-biotin** (57.9 mg, 66.6 %) as a colorless waxy solid: LCMS FA method 1: $R_t$ = 1.101 min, ESI-MS *m/z* 455.4 $(M+2H)^{2+}$, 908.0 $(M+H)^+$, 906.0 $(M-H)^-$; LCMS AA method 1: $R_t$ = 1.376 min, ESI-MS *m/z* 454.6 $(M+2H)^{2+}$, 910.2 $(M+H)^+$, 906.0, 908.2 $(M-H)^-$; $^1$H NMR (600 MHz, DMSO-$d_6$) $\delta_H$ 11.24 (s, 1H), 8.81 (s, 1H), 8.04 (s, 1H), 7.91 (t, *J* = 5.7 Hz, 1H), 7.83 (t, *J* = 5.7 Hz, 1H), 7.78 (dd, *J* = 9.7, 2.1 Hz, 1H), 7.57 (dd, *J* = 8.5, 2.0 Hz, 1H), 7.57 – 7.51 (m, 2H), 7.35 (s, 1H), 6.42 (s, 1H), 6.38 (s, 1H), 4.33 – 4.28 (m, 1H), 4.20 (t, *J* = 6.5 Hz, 2H), 4.13 (dd, *J* = 7.8, 4.4 Hz, 1H), 3.98 (s, 3H), 3.59 (t, *J* = 6.4 Hz, 2H), 3.52 – 3.44 (m, *J* = 3.0 Hz, 12H), 3.38 (t, *J* = 6.0 Hz, 2H), 3.20 – 3.11 (m, 4H), 3.08 (ddd, *J* = 8.7, 6.1, 4.4 Hz, 1H), 2.81 (dd, *J* = 12.5, 5.1 Hz, 1H), 2.60 – 2.52 (m, 1H), 2.31 (t, *J* = 6.4 Hz, 2H), 2.06 (t, *J* = 7.5 Hz, 2H), 1.82 (dt, *J* = 12.3, 6.5 Hz, 2H), 1.65 – 1.55 (m, 2H), 1.55 – 1.39 (m, 4H), 1.36 – 1.22 (m, 2H); $^{13}$C NMR (151 MHz, DMS-$d_6$) $\delta_C$ 172.16, 170.02, 162.73, 158.89, 156.7, 155.99, 150.5, 148.9, 135.82, 130.11, 127.85, 123.92, 120.27, 119.77, 106.73, 103.16, 100.39, 69.63 (4C), 69.45 (2C), 69.05, 68.92, 66.8, 61.01, 59.14, 56.45, 55.33, 39.74, 38.34, 37.86, 36.1, 35.1, 28.05, 27.89, 25.53, 25.48, 25.27.

**N-(4-((4-((4-bromo-2-fluorophenyl)amino)-6-methoxyquinazolin-7-yl)oxy)butyl)-1-(5-((3aS,4S,6aR)-2-oxohexahydro-1H-thieno[3,4-d]imidazol-4-yl)pentanamido)-3,6,9,12,15,18,21,24-octaoxaheptacosan-27-amide 2,2,2-trifluoroacetate (Cp5-PEG8-biotin):**

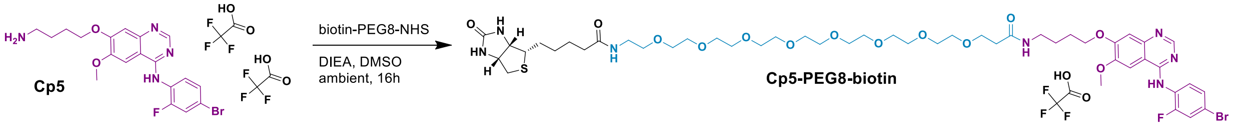


To 7-(4-Aminobutoxy)-N-(4-bromo-2-fluorophenyl)-6-methoxyquinazolin-4-amine bis 2,2,2-trifluoroacetate) (**Cp5**, 14.32 mg, 0.022mmol) and biotin-PEG8-NHS (18.2 mg, 0.024 mmol) dissolved in 1 mL anhydrous DMSO was added DIEA (14 mg, 0.108 mmol). The reaction was shaken at ambient overnight. The reaction was diluted to 3 mL with 90% DMSO/water (v/v) and purified by preparative HPLC TFA method 2. Fractions containing the desired peak were combined and lyophilized to give the desired product **Cp5-PEG8-biotin** (19.4 mg, 74.9 %) as a colorless waxy solid: LCMS FA method 1: $R_t$ = 1.170 min, ESI-MS *m/z* 543.4 $(M+2H)^{2+}$, 1086.4 $(M+H)^+$, 1083.2 $(M-H)^-$; LCMS AA method 1: $R_t$ = 1.414 min, ESI-MS *m/z* 543.8 $(M+2H)^{2+}$, 1086.2 $(M+H)^+$, 1083.2 $(M-H)^-$; $^1$H NMR (600 MHz, DMSO-$d_6$) $\delta_H$ 11.24 (s, 1H), 8.81 (s, 1H), 8.04 (s, 1H), 7.91 (t, *J* = 5.7 Hz, 1H), 7.83 (t, *J* = 5.7 Hz, 1H), 7.78 (dd, *J* = 9.7, 2.1 Hz, 1H), 7.59 – 7.51 (m, 2H), 7.39 – 7.35 (m, 1H), 6.42 (s, 1H), 6.38 (s, 1H), 4.31 (dd, *J* = 7.8, 5.0 Hz, 1H), 4.20 (t, *J* = 6.5 Hz, 2H), 4.13 (dd, *J* = 7.7, 4.4 Hz, 1H), 3.99 (s, 3H), 3.60 (t, *J* = 6.4 Hz, 2H), 3.53 – 3.44 (m, 26H), 3.39 (t, *J* = 5.9 Hz, 2H), 3.21 – 3.06 (m, 6H), 2.81 (dd, *J* = 12.4, 5.1 Hz, 1H), 2.58 (d, *J* = 12.5 Hz, 1H), 2.32 (t, *J* = 6.4 Hz, 2H), 2.06 (t, *J* = 7.4 Hz, 2H), 1.83 (dq, *J* = 12.3, 6.7 Hz, 2H), 1.60 (m, 3H), 1.55 – 1.40 (m, 3H), 1.35 – 1.23 (m, 2H) ; $^{13}$C NMR (151 MHz, DMS-$d_6$) $\delta_C$ 172.16, 170.02, 162.68, 158.87, 156.75, 155.99, 150.41, 148.9, 136.0, 130.07, 127.98, 124.01, 120.3, 119.77, 106.73, 103.04, 100.46, 67.72 (12C), 69.47 (2C), 69.07, 68.93, 66.79, 61.04, 59.11, 56.49, 55.38, 39.76, 38.4, 37.87, 36.17, 35.06, 28.11, 27.95, 25.59, 25.53, 25.18.

**(R)-N-(2-Aminoethyl)-2-chloro-4-((2-((1-hydroxy-3-methylbutan-2-yl)amino)-9-isopropyl-9H-purin-6-yl)amino)benzamide bis-2,2,2-trifluoroacetate (Cp7)** was described (4), but the synthesis was not shown. **Cp7** was prepared using a procedure similar to that described (5).

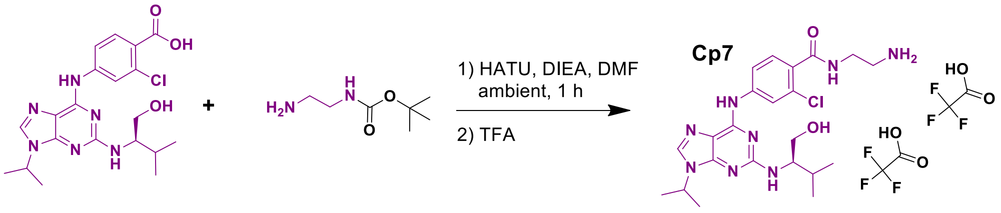


To (R)-2-Chloro-4-((2-((1-hydroxy-3-methylbutan-2-yl)amino)-9-isopropyl-9H-purin-6-yl)amino)benzoic acid (Purvalinol B, 390 mg, 0.901 mmol)) dissolved in 6 mL anhydrous DMF and cooled to < 0 ºC was added HATU (377 mg, 0.991 mmol) and when fully dissolved DIEA (350 mg, 2.70 mmol) and tert-butyl (2-aminoethyl)carbamate (289 mg, 1.80 mmol). The reaction was allowed to warm to ambient and stirred for 2 h. The reaction was diluted to 12 mL in 90% DMSO/water (v/v) and purified by preparative HPLC TFA method 1 in four injections. Fractions containing the desired peak were combined and lyophilized to give the desired **Cp7-Boc** protected intermediate (469.2 mg, 75.6%); LCMS FA method 1: $R_t$ = 1.461 min, ESI-MS *m/z* 575.2 $(M+H)^+$, 573.2 $(M-H)^-$; LCMS AA method 1: $R_t$ = 1.536 min, ESI-MS *m/z* 575.2 $(M+H)^+$, 572.6 $(M-H)^-$; $^1$H NMR (600 MHz, PYRIDINE) $\delta_H$ 10.11 (s, 1H), 9.20 (t, *J* = 5.7 Hz, 1H), 8.08 (s, 1H), 7.88 (d, *J* = 1.1 Hz, 1H), 7.77 (t, *J* = 5.8 Hz, 1H), 7.70 (d, *J* = 17.8 Hz, 1H), 7.08 – 6.91 (m, 1H), 5.69 (s, 2H), 4.60 (hept, *J* = 6.8 Hz, 1H), 4.47 (d, *J* = 6.3 Hz, 1H), 4.15 (qd, *J* = 11.0, 4.9 Hz, 2H), 3.89 (q, *J* = 6.1 Hz, 2H), 3.71 (q, *J* = 6.1 Hz, 2H), 2.37 (h, *J* = 6.8 Hz, 1H), 1.48 (s, 9H), 1.40 (d, *J* = 6.8 Hz, 6H), 1.16 (dd, *J* = 6.8, 4.1 Hz, 6H).

A portion of **Cp7-Boc** (87 mg, 0.126 mmol) was dissolved in 1 mL TFA and immediately evaporated to dryness under a stream of dry nitrogen gas. The residue was dissolved with 3 mL in 90% DMSO/water (v/v) and purified by preparative HPLC TFA method 1 in one injection. Fractions containing the desired peak were combined and lyophilized to give the desired product **Cp7** as a colorless powder (61.5 mg, 69.1%): LCMS FA method 1: $R_t$ = 0.872 min, ESI-MS *m/z* 475.0 $(M+H)^+$, 949.4 $(2M+H)^+$, 473.0 $(M-H)^-$; LCMS AA method 1: $R_t$ = 1.138 min, ESI-MS *m/z* 475.2 $(M+H)^+$, 949.4 $(2M+H)^+$, 473.2 $(M-H)^-$; $^1$H NMR (600 MHz, PYRIDINE) $\delta_H$ 10.15 (s, 1H), 9.67 (t, *J* = 5.7 Hz, 1H), 8.64 – 8.47 (m, 1H), 8.10 (s, 1H), 7.89 (s, 1H), 7.76 (s, 1H), 7.09 – 6.88 (m, 1H), 4.60 (hept, *J* = 6.8 Hz, 1H), 4.47 (d, *J* = 5.9 Hz, 1H), 4.20 – 4.11 (m, 4H), 3.74 (t, *J* = 6.1 Hz, 2H), 1.41 (d, *J* = 6.8 Hz, 6H), 1.17 (dd, *J* = 6.9, 4.2 Hz, 6H).


(4) Médard, G. et al, *J. Proteome Res.*, **2015**, *14*, 1574-1586.
(5) Deane, F.M. et al, *ACS Omega*, **2017**, 2(7), 3828-3838.


**2-chloro-N-(4,20-dioxo-24-((3aS,4S,6aR)-2-oxohexahydro-1H-thieno[3,4-d]imidazol-4-yl)-7,10,13,16-tetraoxa-3,19-diazatetracosyl)-4-((2-(((R)-1-hydroxy-3-methylbutan-2-yl)amino)-9-isopropyl-9H-purin-6-yl)amino)benzamide 2,2,2-trifluoroacetate (Cp7-PEG4-biotin):**

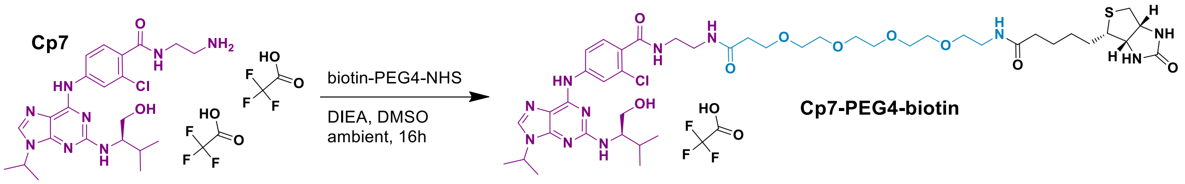


To (R)-N-(2-aminoethyl)-2-chloro-4-((2-((1-hydroxy-3-methylbutan-2-yl)amino)-9-isopropyl-9H-purin-6-yl)amino)benzamide bis-2,2,2-trifluoroacetate **(Cp7,** 90 mg, 127 mmol) and biotin-PEG4-NHS (50 mg, 0.085 mmol) dissolved in 1 mL anhydrous DMSO was added DIEA (60.3 mg, 0.34 mmol). The reaction was shaken at ambient overnight. The reaction was diluted to 3 mL with 90% DMSO/water (v/v) and purified by preparative HPLC TFA method 1. Fractions containing the desired peak were combined and lyophilized to give the desired product **Cp7-PEG4-biotin** (60.3 mg, 67.0 %) as a colorless waxy solid: LCMS FA method 1: $R_t$ = 1.212 min, ESI-MS *m/z* 474.6 $(M+2H)^{2+}$, 958.2 $(M+H)^+$, 946.4 $(M-H)^-$; LCMS AA method 1: $R_t$ = 1.297 min, ESI-MS *m/z* 474.8 $(M+2H)^{2+}$, 948.2 $(M+H)^+$, 946.6 $(M-H)^-$; $^1$H NMR (600 MHz, DMSO-*d*$_6$) $\delta_H$ 10.11 (s, 1H), 8.35 (d, J = 3.3 Hz, 1H), 8.29 (t, J = 5.6 Hz, 1H), 8.21 (s, 1H), 7.93 (t, *J* = 5.7 Hz, 1H), 7.89 (t, *J* = 5.7 Hz, 1H), 7.82 (t, *J* = 5.7 Hz, 1H), 7.43 (d, *J* = 8.4 Hz, 1H), 6.78 (s, 1H), 6.41 (s, 2H), 4.65 (hept, *J* = 6.6 Hz, 1H), 4.37 - 4.26 (m, 1H)), 4.12 (dd, *J* = 7.8, 4.4 Hz, 1H), 3.87 (m, 1H), 3.63 – 3.53 (m, 2H), 3.53-3.44 (m, 12H), 3.31-3.18 (m, 6H), 3.08 (ddd, *J* = 8.l6, 6.1, 4.4 Hz, 1H), 2.81 (dd. *J* = 12.4, 5.1 Hz, 1H), 2.63 – 2.53 (m, 1H), 2.5 (m, 3H), 2.33 (t, *J* = 6.5 Hz, 2H), 2.06 (t, *J* = 7.4 Hz, 2H), 1.97 (h, *J* = 6.8 Hz, 1H), 1.66-1.57 (m, 1H), 1.56-1.39 (m, 9H), 1.38 – 1.19 (m, 2H), 0.95 (t, *J* = 6.4 Hz, 6H); $^{13}$C NMR (151 MHz, DMS-*d*$_6$) $\delta_C$ 173.94, 172.17, 170.37, 166.15, 162.73, 158.05, 150.63, 149.38, 141.3, 136.64, 130.23, 130.09, 129.32, 120.44, 118.24, 117.28, 114.85, 112.58, 111.44, 95.26, 89.11, 80.11, 69.7 (2H), 69.64, 69.60, 69.48, 69.47, 69.07, 66.67, 61.00, 60.86, 59.16, 58.56, 55.33, 46.87, 39.82, 38.92, 38.39, 38.07, 36.12, 34.99, 28.87, 28.11, 28.00, 25.12, 21.77, 21.62, 19.37, 18.86.

**2-chloro-N-(4,32-dioxo-36-((3aS,4S,6aR)-2-oxohexahydro-1H-thieno[3,4-d]imidazol-4-yl)-7,10,13,16,19,22,25,28-octaoxa-3,31-diazahexatriacontyl)-4-((2-(((R)-1-hydroxy-3-methylbutan-2-yl)amino)-9-isopropyl-9H-purin-6-yl)amino)benzamide 2,2,2-trifluoroacetate (Cp7-PEG8-biotin):**

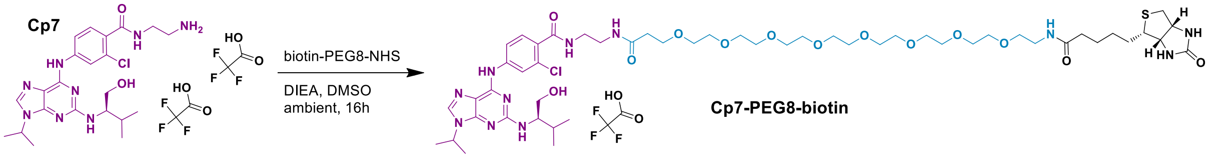


To (R)-N-(2-aminoethyl)-2-chloro-4-((2-((1-hydroxy-3-methylbutan-2-yl)amino)-9-isopropyl-9H-purin-6-yl)amino)benzamide bis-2,2,2-trifluoroacetate **(Cp7,** 19 mg, 0.027 mmol) and biotin-PEG8-NHS (24.8 mg, 0.032 mmol) dissolved in 1 mL anhydrous DMSO was added DIEA (14.8 mg, 0.115 mmol). The reaction was shaken at ambient overnight. The reaction was diluted to 3 mL with 90% DMSO/water (v/v) and purified by preparative HPLC TFA method 2. Fractions containing the desired peak were combined and lyophilized to give the desired product **Cp7-PEG8-biotin** (12.5 mg, 37.3 %) as a colorless waxy solid: LCMS FA method 1: $R_t$ = 1.254 min, ESI-MS *m/z* 562.6 $(M+2H)^{2+}$, 1125.4 $(M+H)^+$, 1123.2 $(M-H)^-$; LCMS AA method 1: $R_t$ = 1.334 min, ESI-MS *m/z* 562.6 $(M+2H)^{2+}$, 1124.8 $(M+H)^+$, 1123.2 $(M-H)^-$; $^1$H NMR (600 MHz, DMSO) $\delta_H$ 10.10 (s, 1H), 8.33 (s, 1H), 8.29 (t, *J* = 5.6 Hz, 1H), 8.21 (s, 1H), 7.93 (t, *J* = 5.7 Hz, 1H), 7.86 (s, 1H), 7.82 (t, *J* = 5.7 Hz, 1H), 7.43 (d, *J* = 8.4 Hz, 1H), 6.75 (s, 1H), 6.40 (s, 2H), 4.64 (h, *J* = 6.8 Hz, 1H), 4.30 (dd, *J* = 7.7, 4.9 Hz, 1H), 4.13 (dd, *J* = 7.7, 4.4 Hz, 1H), 3.86 (q, *J* = 5.7 Hz, 1H), 3.60, (m, 2H), 3.53 – 3.44 (m, 24H), 3.56 – 3.57 (m, 2H), 3.38 (t, *J* = 5.9 Hz, 2H), 3.31 – 3.20 (m, 4H), 3.09 (ddd, *J* = 8.6, 6.1, 4.4 Hz, 1H), 2.80, (m, 1H), 2.58 (d, *J* = 12.5 Hz, 1H), 2.50 (p, *J* = 1.8 Hz, 3H), 2.32 (t, *J* = 6.5 Hz, 2H), 2.06 (t, *J* = 7.4 Hz, 2H), 1.97 (h, *J* = 6.8 Hz, 1H), 1.67 – 1.57 (m, 1H), 1.57 – 1.40 (m, 8H), 1.38

– 1.19 (m, 2H), 0.94 (t, $J$ = 6.5 Hz, 6H); $^{13}$C NMR (151 MHz, DMS-$d_6$) $\delta_C$ 174.63, 172.24, 170.45, 165.97, 162.64, 157.99, 150.39, 149.34, 141.36, 136.95, 130.04, 129.94, 129.14, 120.43, 118.35, 117.28, 115.03, 112.60, 111.47, 94.31, 89.00, 80.10, 69.52 (10C), 69.47, 69.41, 69.31, 69.30, 69.13, 66.79, 61.04, 60.82, 59.07, 58.28, 55.41, 46.95, 39.81, 38.90, 38.22, 37.95, 35.98, 34.85, 28.68, 28.03, 27.93, 25.21, 21.49 (2C), 19.01 (2C).

**tert-Butyl (2-(2-(2-((N-(4-((2-(4-amino-1,2,5-oxadiazol-3-yl)-1-ethyl-1H-imidazo[4,5-c]pyridin-7-yl)oxy)butyl)-2-nitrophenyl)sulfonamido)ethoxy)ethoxy)ethyl)carbamate (Cp15-Ns, Boc)** was synthesized by WuXi Apptec, Shanghai, China as described (6)(7).

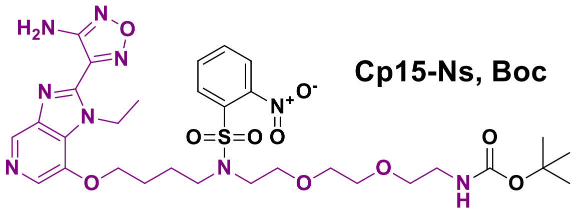


LCMS FA method 1: $R_t$ = 1.472 min, ESI-MS *m/z* 734.2 (M+H)$^+$; LCMS AA method 1: $R_t$ = 1.828 min, ESI-MS *m/z* 734.2 (M+H)$^+$; $^1$H NMR (400 MHz, CDCl3) $\delta_H$ 8.80 (s, 1H), 8.07-8.05 (m,2H), 7.70 7.64 (m, 3H), 5.91 (s, 2H), 4.98-4.95 (m, 3H), 4.25 (t, J = 5.7 Hz, 2H), 3.64-3.61 (m, 2H), 3.56-3.48 (m,10H), 3.30-3.28 (m, 2H), 1.89-1.85 (br, 4H), 1.50 (t, J = 7.0 Hz, 3H), 1.43 (s, 9H).

(6) Heerding, D.A. et al, *J. Med. Chem.*, **2008**, *51(18),* 5663-5679.
(7) Pachl, F. et al, J. Proteome. Res. **2013**, *12*, 3792-3800.

**N-(4-((2-(4-Amino-1,2,5-oxadiazol-3-yl)-1-ethyl-1H-imidazo[4,5-c]pyridin-7-yl)oxy)butyl)-N-(2-(2-(2-aminoethoxy)ethoxy)ethyl)-2-nitrobenzenesulfonamide bis(2,2,2-trifluoroacetate) (Cp15-Ns):**

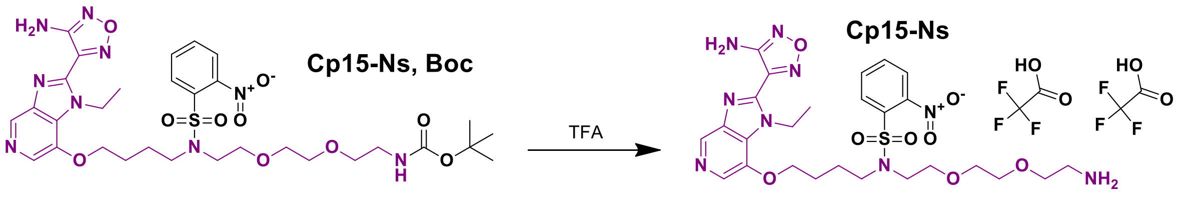


tert-Butyl (2-(2-(2-((N-(4-((2-(4-amino-1,2,5-oxadiazol-3-yl)-1-ethyl-1H-imidazo[4,5-c]pyridin-7-yl)oxy)butyl)-2-nitrophenyl)sulfonamido)ethoxy)ethoxy)ethyl)carbamate **(Cp15-Ns, Boc**, 100 mg, 0.136 mmol) was dissolved in 1 mL TFA and immediately evaporated to dryness under a stream of dry nitrogen gas to give the desired product **Cp15-Ns** as a white solid (117 mg, 100%): LCMS FA method 1: $R_t$ = 0.928 min, ESI-MS *m/z* 317.8 (M+2H)$^{2+}$, 634.2 (M+H)$^+$, no negative ion; LCMS AA method 1: $R_t$ = 1.456 min, ESI-MS *m/z* 634.2 (M+H)$^+$, 632.2 (M-H)$^-$.

**4-(7-(4-((2-(2-(2-Aminoethoxy)ethoxy)ethyl)amino)butoxy)-1-ethyl-1H-imidazo[4,5-c]pyridin-2-yl)-1,2,5-oxadiazol-3-amine tris(2,2,2-trifluoroacetate) (Cp15):**

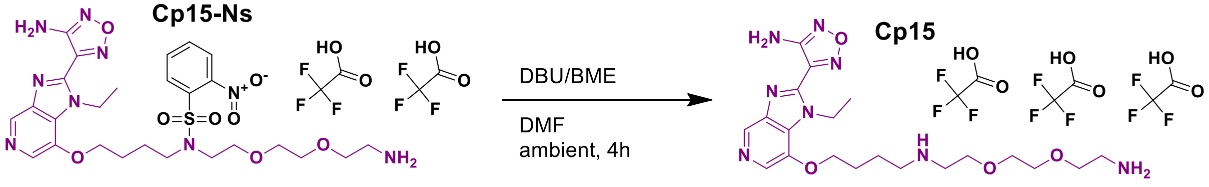


To N-(4-((2-(4-Amino-1,2,5-oxadiazol-3-yl)-1-ethyl-1H-imidazo[4,5-c]pyridin-7-yl)oxy)butyl)-N-(2-(2-(2-aminoethoxy)ethoxy)ethyl)-2-nitrobenzenesulfonamide bis(2,2,2-trifluoroacetate) (**Cp15-Ns**, 30 mg, 0.035 mmol) dissolved in 1 mL anhydrous DMF was added 1,8-diazabicyclo[5.4.0]undec-7-ene (DBU, 101 mg, 0.663 mmol) and 2-mercaptoethanol (BME, 55 mg, 0.710 mmol). The reaction was shaken at ambient for during which time the solution became yellow. After 2 h. the reaction was diluted to 3 mL in 90% DMSO/water (v/v) and purified by preparative HPLC TFA method 1 in one injection. Fractions containing the desired peak were combined and lyophilized to give the desired product **Cp15** as a colorless powder (18.6 mg, 67.6%): LCMS FA method 1: $R_t = 0.146$ min, ESI-MS *m/z* 449.2 $(M+H)^+$; LCMS AA method 1 (broad with tailing): $R_t = 0.947$ min, ESI-MS *m/z* 449.2 $(M+H)^+$.

**N-(2-(2-(2-((N-(4-((2-(4-Amino-1,2,5-oxadiazol-3-yl)-1-ethyl-1H-imidazo[4,5-c]pyridin-7-yl)oxy)butyl)-2-nitrophenyl)sulfonamido)ethoxy)ethoxy)ethyl)-1-(5-((3aS,4S,6aR)-2-oxohexahydro-1H-thieno[3,4-d]imidazol-4-yl)pentanamido)-3,6,9,12-tetraoxapentadecan-15-amide 2,2,2-trifluoroacetate (Cp15-Ns-PEG4-biotin):**

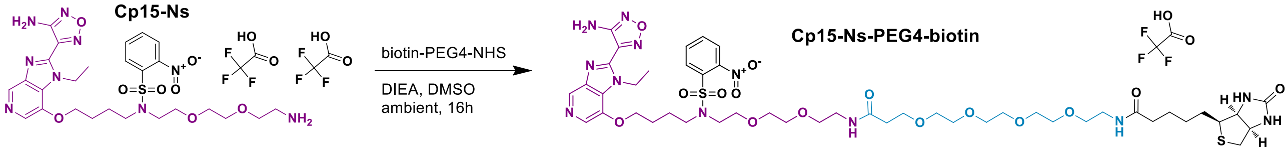


To N-(4-((2-(4-Amino-1,2,5-oxadiazol-3-yl)-1-ethyl-1H-imidazo[4,5-c]pyridin-7-yl)oxy)butyl)-N-(2-(2-(2-aminoethoxy)ethoxy)ethyl)-2-nitrobenzenesulfonamide bis(2,2,2-trifluoroacetate) (**Cp15-Ns**, 60 mg, 0.07 mmol) and biotin-PEG4-NHS (50 mg, 0.085 mmol) dissolved in 1 mL anhydrous DMSO was added DIEA (60 mg, 0.464 mmol). The reaction was shaken at ambient overnight. The reaction was diluted to 3 mL with 90% DMSO/water (v/v) and purified by preparative HPLC TFA method 2. Fractions containing the desired peak were combined and lyophilized to give the desired product **Cp15-Ns-PEG4-biotin** (35.1 mg, 41.3 %) as a colorless waxy solid: LCMS FA method 1: $R_t = 1.228$ min, ESI-MS *m/z* 554.2 $(M+2H)^{2+}$, 1108.4 $(M+H)^+$, 1105.4 $(M-H)^-$; LCMS AA method 1: $R_t = 1.499$ min, ESI-MS *m/z* 554.4 $(M+2H)^{2+}$, 1107.2 $(M+H)^+$, 1105.4 $(M-H)^-$.

**Method 1 - N-(2-(2-(2-((4-((2-(4-Amino-1,2,5-oxadiazol-3-yl)-1-ethyl-1H-imidazo[4,5-c]pyridin-7-yl)oxy)butyl)amino)ethoxy)ethoxy)ethyl)-1-(5-((3aS,4S,6aR)-2-oxohexahydro-1H-thieno[3,4-d]imidazol-4-yl)pentanamido)-3,6,9,12-tetraoxapentadecan-15-amide bis-2,2,2-trifluoroacetate (Cp15-PEG4-biotin):**

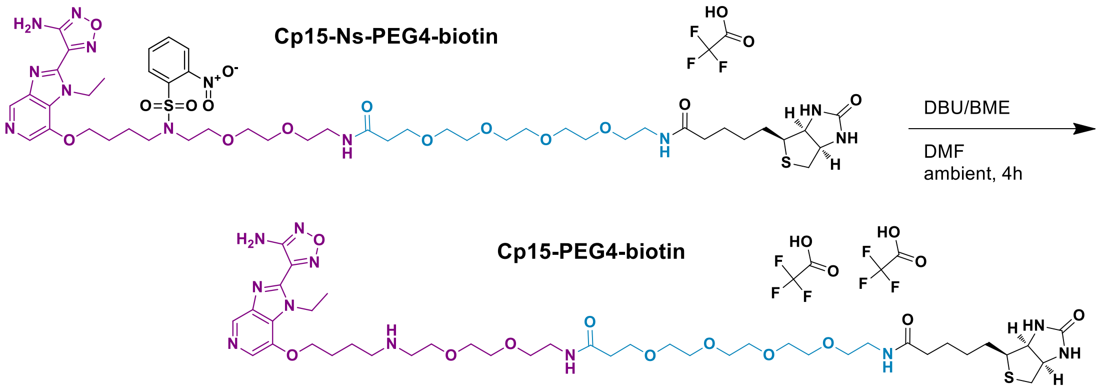


N-(2-(2-(2-((N-(4-((2-(4-Amino-1,2,5-oxadiazol-3-yl)-1-ethyl-1H-imidazo[4,5-c]pyridin-7-yl)oxy)butyl)-2-nitrophenyl)sulfonamido)ethoxy)ethoxy)ethyl)-1-(5-((3aS,4S,6aR)-2-oxohexahydro-1H-thieno[3,4-d]imidazol-4-yl)pentanamido)-3,6,9,12-tetraoxapentadecan-15-amide 2,2,2-trifluoroacetate **(Cp15-Ns-PEG4-biotin**, 11.6 mg, 0.0105 mmol) was dissolved in 0.5 mL anhydrous DMF. 2-Mercaptoethanol (BME, 20 mg, 0.256 mmol) and 1,8-diazabicyclo[5.4.0]undec-7-ene (DBU, 40 mg, 0.263 mmol) were added and the reaction shaken at ambient for 4h. The reaction was diluted to 3 mL with 90% DMSO/water (v/v) and purified by preparative HPLC TFA method 2. Fractions containing the desired peak were combined and lyophilized to give the desired product **Cp15-PEG4-biotin** (10.2 mg, 84.6 %) as a colorless waxy solid: LCMS FA method 1: $R_t$ = 0.814 min, ESI-MS *m/z* 462.4 $(M+2H)^{2+}$, 922.6 $(M+H)^+$, 920.2 $(M-H)^-$; LCMS AA method 1: $R_t$ = 1.143 min, ESI-MS *m/z* 461.8 $(M+2H)^{2+}$, 922.2 $(M+H)^+$, 921.2 $(M-H)^-$.

**Method 2 - N-(2-(2-(2-((4-((2-(4-Amino-1,2,5-oxadiazol-3-yl)-1-ethyl-1H-imidazo[4,5-c]pyridin-7-yl)oxy)butyl)amino)ethoxy)ethoxy)ethyl)-1-(5-((3aS,4S,6aR)-2-oxohexahydro-1H-thieno[3,4-d]imidazol-4-yl)pentanamido)-3,6,9,12-tetraoxapentadecan-15-amide bis-2,2,2-trifluoroacetate (Cp15-PEG4-biotin):**

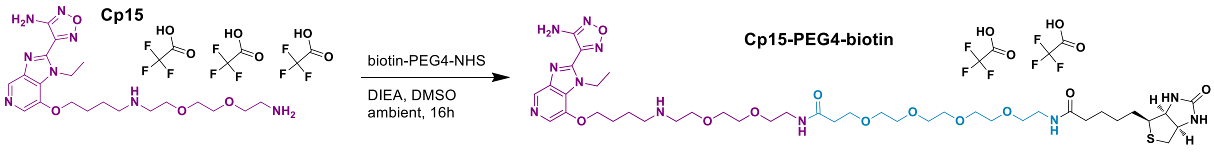


To 4-(7-(4-((2-(2-(2-Aminoethoxy)ethoxy)ethyl)amino)butoxy)-1-ethyl-1H-imidazo[4,5-c]pyridin-2-yl)-1,2,5-oxadiazol-3-amine tris(2,2,2-trifluoroacetate) (**Cp15**, 72 mg, 0.091 mmol) and biotin-PEG4-NHS (50 mg, 0.085 mmol) dissolved in 1 mL anhydrous DMSO was added DIEA (44 mg, 0.34 mmol). The reaction was shaken at ambient overnight. The reaction was diluted to 3 mL with 90% DMSO/water (v/v) and purified by preparative HPLC TFA method 1. Fractions containing the desired peak were combined and lyophilized to give the desired product **Cp15-PEG4-biotin** (71.4 mg, 72.6 %) as a colorless waxy solid: LCMS TFA method 1: $R_t$ = 0.60 min, ESI-MS *m/z* 449.38 $(M+2H)^{2+}$; $^1$H NMR (400 MHz, DMSO) $\delta_H$ 8.55 (s, 1H), 7.89 – 7.75 (m, 1H), 6.92 (s, 1H), 4.84 (q, *J* = 7.0 Hz, 1H), 4.38 – 4.23 (m, 1H), 3.64 (t, *J* = 5.1 Hz, 1H), 3.55 (dt, *J* = 8.2, 5.4 Hz, 8H), 3.53 – 3.31 (m, 6H), 3.15 (tt, *J* = 11.4, 5.7 Hz, 2H), 3.12 – 2.97 (m, 1H), 2.27 (t, *J* = 6.5 Hz, 1H), 2.02 (t, *J* = 7.4 Hz, 1H), 1.94 – 1.77 (m, 2H), 1.45 (q, *J* = 6.9 Hz, 2H), 1.26 (tt, *J* = 14.5, 6.5 Hz, 1H).

**N-(2-(2-(2-((N-(4-((2-(4-Amino-1,2,5-oxadiazol-3-yl)-1-ethyl-1H-imidazo[4,5-c]pyridin-7-yl)oxy)butyl)-2-nitrophenyl)sulfonamido)ethoxy)ethoxy)ethyl)-1-(5-((3aS,4S,6aR)-2-oxohexahydro-1H-thieno[3,4-d]imidazol-4-yl)pentanamido)-3,6,9,12,15,18,21,24-octaoxaheptacosan-27-amide 2,2,2-trifluoroacetate (Cp15-Ns-PEG8-biotin):**

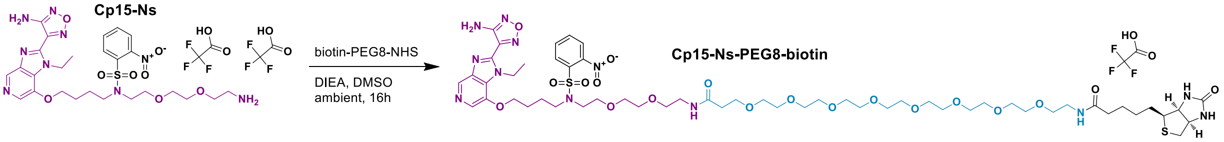


To N-(4-((2-(4-Amino-1,2,5-oxadiazol-3-yl)-1-ethyl-1H-imidazo[4,5-c]pyridin-7-yl)oxy)butyl)-N-(2-(2-(2-aminoethoxy)ethoxy)ethyl)-2-nitrobenzenesulfonamide bis(2,2,2-trifluoroacetate) **(Cp15-Ns**, 47.2 mg, 0.055 mmol) and biotin-PEG8-NHS (41.9 mg, 0.055 mmol) dissolved in 1 mL anhydrous DMSO was added DIEA (48 mg, 0.274 mmol). The reaction was shaken at ambient overnight. The reaction was diluted to 3 mL with 90% DMSO/water (v/v) and purified by preparative HPLC TFA method 2. Fractions containing the desired peak were combined and lyophilized to give the desired product **Cp15-Ns-PEG8-biotin** (52.8 mg, 68.6 %) as a colorless waxy solid: LCMS FA method 1: $R_t$ = 1.260 min, ESI-MS *m/z* 642.2 $(M+2H)^{2+}$, 1283.2 $(M+H)^+$, 1282.2 $(M-H)^-$; LCMS AA method 1: $R_t$ = 1.525 min, ESI-MS *m/z* 642.0 $(M+2H)^{2+}$, 1281.4 $(M-H)^-$.

**N-(2-(2-(2-((4-((2-(4-Amino-1,2,5-oxadiazol-3-yl)-1-ethyl-1H-imidazo[4,5-c]pyridin-7-yl)oxy)butyl)amino)ethoxy)ethoxy)ethyl)-1-(5-((3aS,4S,6aR)-2-oxohexahydro-1H-thieno[3,4-d]imidazol-4-yl)pentanamido)-3,6,9,12,15,18,21,24-octaoxaheptacosan-27-amide bis-2,2,2-trifluoroacetate (Cp15-PEG8-biotin):**

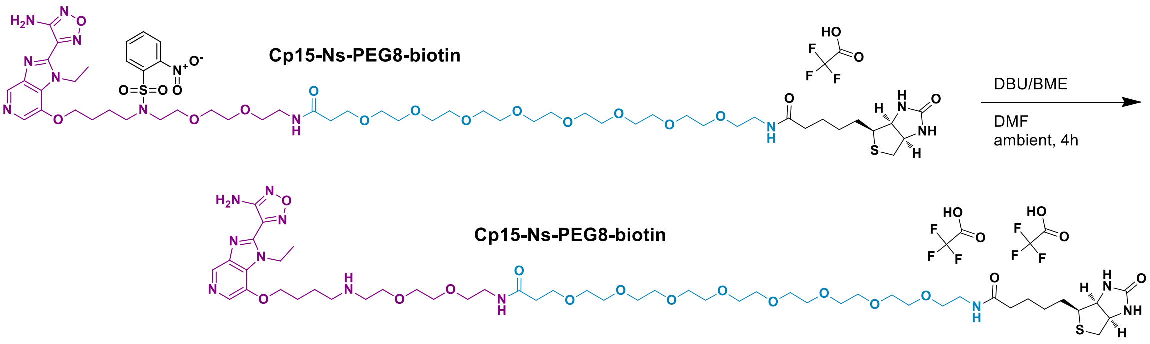


N-(2-(2-(2-((N-(4-((2-(4-Amino-1,2,5-oxadiazol-3-yl)-1-ethyl-1H-imidazo[4,5-c]pyridin-7-yl)oxy)butyl)-2-nitrophenyl)sulfonamido)ethoxy)ethoxy)ethyl)-1-(5-((3aS,4S,6aR)-2-oxohexahydro-1H-thieno[3,4-d]imidazol-4-yl)pentanamido)-3,6,9,12,15,18,21,24-octaoxaheptacosan-27-amide 2,2,2-trifluoroacetate **(Cp15-Ns-PEG8-biotin**, 18.8 mg, 0.013 mmol) was dissolved in 0.5 mL anhydrous DMF. 2-Mercaptoethanol (BME, 25 mg, 0.320 mmol) and 1,8-diazabicyclo[5.4.0]undec-7-ene (DBU, 50 mg, 0.328 mmol) were added and the reaction shaken at ambient for 4h. The reaction was diluted to 3 mL with 90% DMSO/water (v/v) and purified by preparative HPLC TFA method 2. Fractions containing the desired peak were combined and lyophilized to give the desired product **Cp15-PEG8-biotin** (4.6 mg, 25.8 %) as a colorless waxy solid: LCMS TFA method 1: $R_t$ = 0.60 min, ESI-MS *m/z* 550.0 $(M+2H)^{2+}$; LCMS AA method 1: $R_t$ = 1.19 min, ESI-MS *m/z* 549.8 $(M+2H)^{2+}$, 1096.4 $(M-H)^-$; $^1$H NMR (600 MHz, DMSO) $\delta_H$ 9.04 (s, 1H), 8.73 (s, 2H), 8.35 (s, 1H), 7.91 (q, *J* = 4.4 Hz, 1H), 7.83 (t, *J* = 5.7 Hz, 1H), 6.95 (s, 2H), 6.40

(s, 1H), 6.35 (s, 1H), 4.88 (q, $J$ = 7.1 Hz, 2H), 4.37 (t, $J$ = 6.0 Hz, 2H), 4.33 – 4.28 (m, 1H), 4.13 (dd, $J$ = 7.8, 4.4 Hz, 1H), 3.69 (t, $J$ = 5.2 Hz, 2H), 3.61 – 3.57 (m, 2H), 3.59 – 3.53 (m, 4H), 3.53 – 3.47 (m, 24H), 3.50 – 3.43 (m, 4H), 3.40 (dt, $J$ = 9.3, 6.0 Hz, 4H), 3.19 (dq, $J$ = 13.7, 5.9 Hz, 4H), 3.16 – 3.02 (m, 5H), 2.82 (dd, $J$ = 12.4, 5.1 Hz, 1H), 2.58 (d, $J$ = 12.4 Hz, 1H), 2.34 – 2.25 (m, 2H), 2.06 (t, $J$ = 7.4 Hz, 2H), 1.94 (dq, $J$ = 11.4, 6.4 Hz, 2H), 1.86 (ddd, $J$ = 15.3, 8.9, 5.8 Hz, 2H), 1.61 (ddt, $J$ = 13.6, 9.9, 6.1 Hz, 1H), 1.56 – 1.40 (m, 5H), 1.30 (dddt, $J$ = 15.4, 9.4, 6.0, 2.9 Hz, 2H); $^{13}$C NMR (151 MHz, DMSO) $\delta_C$ 172.09, 170.19, 162.65, 156.25, 143.71, 143.62, 139.92, 137.60, 133.72, 131.17, 122.61, 69.7 (11C), 69.64, 69.59, 69.50, 69.44, 69.29, 69.09, 69.08, 68.95, 66.69, 65.56, 60.98, 59.15, 55.34, 46.54, 46.19, 43.17, 39.76, 38.38, 38.37, 35.95, 35.03, 28.12, 27.96, 25.69, 25.18, 22.14, 16.03.

**2-((2-((4-(aminomethyl)phenyl)amino)-5-chloropyrimidin-4-yl)amino)-N-methylbenzenesulfonamide (Cp19)** was synthesized by WuXi Apptec, Shanghai, China as described (8).

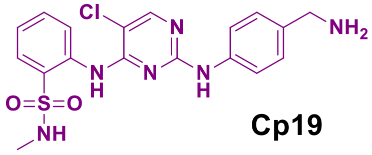


**Cp19**

LCMS FA method 1: $R_t$ = 0.862 min, ESI-MS *m/z* 210.0 $(M+2H)^{2+}$, 419.0 $(M+H)^+$, 417.0 $(M-H)^-$; LCMS AA method 1: $R_t$ = 1.170 min, ESI-MS *m/z* 419.0 $(M+H)^+$, 417.0 $(M-H)^-$; $^1$H NMR (400 MHz, DMSO) $\delta_H$ 9.43 (s, 1H), 8.54 (d, J = 8.2 Hz, 1H), 8.25 (s, 1H), 7.82 (dd, J = 8.0, 1.5 Hz, 1H), 7.70 - 7.61 (m, 1H), 7.56 (d, J = 8.5 Hz, 2H), 7.36 - 7.26 (m, 1H), 7.20 (d, J = 8.5 Hz, 2H), 3.66 (s, 2H), 2.43 (s, 3H).


(8) Chan, S., et al, *Bioorg. Med. Chem. Lett.*, **2015**, 25(19), 4277-4281.


***Tert*-butyl (2-(2-(2-((4-((5-chloro-4-((2-(N-methylsulfamoyl)phenyl)amino)pyrimidin-2-yl)amino)benzyl)amino)-2-oxoethoxy)ethoxy)ethyl)carbamate 2,2,2-trifluoroacetate (Cp19-PEG2-NHBoc):**

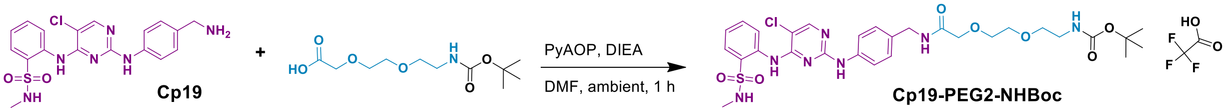


2-((2-((4-(Aminomethyl)phenyl)amino)-5-chloropyrimidin-4-yl)amino)-N-methylbenzenesulfonamide (**Cp19**, 103.5 mg, 0.247 mmol) and 2,2-dimethyl-4-oxo-3,8,11-trioxa-5-azatridecan-13-oic acid (78 mg, 0.296 mmol) were combined in 1 mL anhydrous DMF. ((3H-[1,2,3]Triazolo[4,5-b]pyridin-3-yl)oxy)tri(pyrrolidin-1-yl)phosphonium hexafluorophosphate(V) (PyAOP, 155 mg, 0.296 mmol) and N-ethyl-N-isopropylpropan-2-amine (214 µl, 1.235 mmol) were added and the reaction shaken at ambient for 1h. The reaction was diluted to 3 mL with 90% DMSO/water (v/v) and purified by preparative HPLC TFA method 1. Fractions containing the desired peak were combined and lyophilized to give the desired product **Cp19-PEG2-NHBoc** (160 mg, 83.3 %) as a colorless solid: LCMS TFA method 1: $R_t$ = 0.88 min, ESI-MS *m/z* 664.12 $(M+H)^+$; $^1$H NMR (400 MHz, DMSO) $\delta_H$ 9.43 (s, 1H), 9.26 (s, 1H), 8.50 (d, $J$ = 8.3 Hz, 1H), 8.23 (s, 1H), 8.11 (t, $J$ = 6.2 Hz, 1H), 7.79 (dd, $J$ = 8.0, 1.6 Hz, 1H), 7.74 (s, 1H), 7.60 (ddd, $J$ = 8.7, 7.3, 1.6 Hz, 1H), 7.52 (d, $J$ = 8.5 Hz, 2H), 7.28 (ddd, $J$ = 8.2, 7.4, 1.2 Hz, 1H), 7.15 – 7.08 (m, 2H), 6.71 (d, $J$ = 5.7 Hz, 1H), 4.22 (d, $J$ = 6.2 Hz, 2H), 3.90 (s, 2H), 3.56 (dd, $J$ = 6.2, 3.6 Hz, 2H), 3.51 (dd, $J$ = 6.2, 3.6 Hz, 2H), 3.01 (q, $J$ = 6.0 Hz, 2H), 2.40 (s, 3H), 1.32 (s, 9H).

**2-(2-(2-aminoethoxy)ethoxy)-N-(4-((5-chloro-4-((2-(N-methylsulfamoyl)phenyl)amino)pyrimidin-2-yl)amino)benzyl)acetamide bis-(2,2,2-trifluoroacetate) (Cp19-PEG2-NH$_2$):**

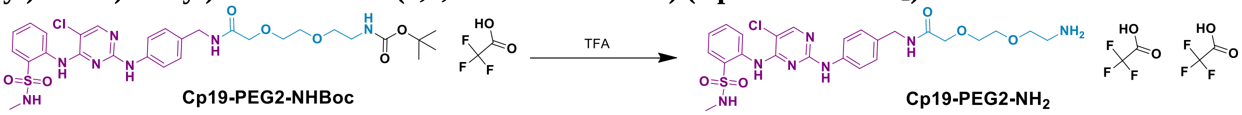


*Tert*-butyl (2-(2-(2-((4-((5-chloro-4-((2-(N-methylsulfamoyl)phenyl)amino)pyrimidin-2-yl)amino)benzyl)amino)-2-oxoethoxy)ethoxy)ethyl)carbamate 2,2,2-trifluoroacetate (Cp19-PEG2-NHBoc, 145 mg, 0.186 mmol) was dissolved in 1 mL trifluoroacetic acid and immediately evaporated to dryness under a stream of dry nitrogen gas to give the desired product **Cp19-PEG2-NH$_2$** as a colorless glassy solid (148 mg, 100 %): LCMS TFA method 1: $R_t$ = 0.69 min, ESI-MS *m/z* 564.22 $(M+H)^+$.

**N-(2-(2-(2-((4-((5-chloro-4-((2-(N-methylsulfamoyl)phenyl)amino)pyrimidin-2-yl)amino)benzyl)amino)-2-oxoethoxy)ethoxy)ethyl)-5-((3aS,4S,6aR)-2-oxohexahydro-1H-thieno[3,4-d]imidazol-4-yl)pentanamide 2,2,2-trifluoroacetate (Cp19-PEG2-biotin):**

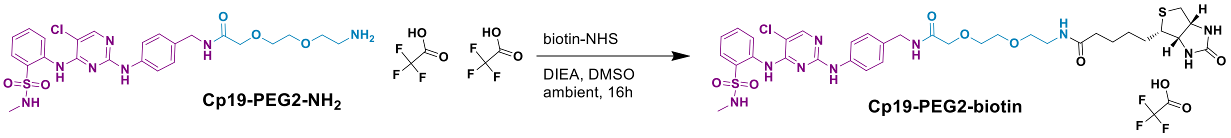


To 2-(2-(2-aminoethoxy)ethoxy)-N-(4-((5-chloro-4-((2-(N-methylsulfamoyl)phenyl)amino)pyrimidin-2-yl)amino)benzyl)acetamide bis-(2,2,2-trifluoroacetate) (**Cp19-PEG2-NH$_2$**, 31.7 mg, 0.04 mmol) and biotin-PEG4-NHS (50 mg, 0.085 mmol) in 1 mL anhydrous dimethylsulfoxide was added N-ethyl-N-isopropylpropan-2-amine (59.3 µl, 0.34 mmol) and the reaction shaken at ambient for 16 h. The reaction was diluted to 3 mL with 90% DMSO/water (v/v) and purified by preparative HPLC TFA method 1. Fractions containing the desired peak were combined and lyophilized to give the desired product **Cp19-PEG2-biotin** (34.0 mg, 93.9 %) as a colorless waxy solid: LCMS FA method 1: $R_t$ = 1.27 min, ESI-MS *m/z* 790.0 $(M+H)^+$. LCMS AA method 1: $R_t$ = 1.35 min, ESI-MS *m/z* 790.0 $(M+H)^+$. $^1$H NMR (499 MHz, DMSO) δ 9.54 (s, 1H), 9.36 (s, 1H), 8.50 (d, *J* = 8.3 Hz, 1H), 8.28 (s, 1H), 8.16 (t, *J* = 6.2 Hz, 1H), 7.88 – 7.70 (m, 3H), 7.64 (ddd, *J* = 8.6, 7.4, 1.6 Hz, 1H), 7.54 (d, *J* = 8.2 Hz, 2H), 7.33 (ddd, *J* = 8.3, 7.4, 1.2 Hz, 1H), 7.21 – 7.07 (m, 2H), 6.40 (s, 1H), 4.31 – 4.25 (m, 2H), 4.11 (dd, *J* = 7.8, 4.4 Hz, 1H), 3.94 (s, 2H), 3.66 – 3.49 (m, 4H), 3.40 (t, *J* = 5.9 Hz, 2H), 3.18 (t, *J* = 5.8 Hz, 2H), 3.07 (ddd, *J* = 8.6, 6.2, 4.4 Hz, 1H), 2.80 (dd, *J* = 12.5, 5.1 Hz, 1H), 2.57 (d, *J* = 12.4 Hz, 1H), 2.50 (p, *J* = 1.9 Hz, 3H), 2.44 (d, *J* = 4.8 Hz, 3H), 2.05 (t, *J* = 7.4 Hz, 2H), 1.59 (ddt, *J* = 12.1, 9.7, 4.7 Hz, 1H), 1.47 (dddd, *J* = 19.2, 14.4, 6.9, 3.8 Hz, 2H), 1.33 – 1.13 (m, 2H).

**N-(4-((5-chloro-4-((2-(N-methylsulfamoyl)phenyl)amino)pyrimidin-2-yl)amino)benzyl)-1-(5-((3aS,4S,6aR)-2-oxohexahydro-1H-thieno[3,4-d]imidazol-4-yl)pentanamido)-3,6,9,12-tetraoxapentadecan-15-amide 2,2,2-trifluoroacetate (Cp19-PEG4-biotin):**

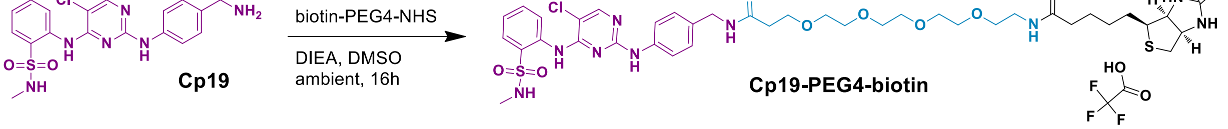

To 2-((2-((4-(Aminomethyl)phenyl)amino)-5-chloropyrimidin-4-yl)amino)-N-methylbenzenesulfonamide (**Cp19**, 53.4 mg, 0.127 mmol) and biotin-PEG4-NHS (16.4 mg, 0.028 mmol) in 1 mL anhydrous dimethylsulfoxide was added N-ethyl-N-isopropylpropan-2-amine (20 µl, 0.115 mmol) and the reaction shaken at ambient for 16 h. The reaction was diluted to 3 mL with 90% DMSO/water (v/v) and purified by preparative HPLC TFA method 2. Fractions containing the desired peak were combined and lyophilized to give the desired product **Cp19-PEG4-biotin** (53.3 mg, 62.7 %) as a colorless waxy solid: LCMS FA method 1: $R_t$ = 1.31 min, ESI-MS *m/z* 446.6 $(M+2H)^{2+}$, 892.2 $(M+H)^+$, 890.2 $(M-H)^-$; LCMS AA method 1: $R_t$ = 1.36 min, ESI-MS *m/z* 447.0 $(M+2H)^{2+}$, 892.0 $(M+H)^+$, 890.0 $(M-H)^-$; $^1$H NMR (600 MHz, DMSO) $\delta_H$ 9.61 (s, 1H), 9.42 (s, 1H), 8.49 – 8.45 (m, 1H), 8.29 (t, *J* = 6.0 Hz, 1H), 8.29 (s, 1H), 7.85 – 7.79 (m, 2H), 7.76 (q, *J* = 4.9 Hz, 1H), 7.65 (ddd, *J* = 8.6, 7.3, 1.6 Hz, 1H), 7.52 (d, *J* = 8.1 Hz, 2H), 7.34 (td, *J* = 7.6, 1.2 Hz, 1H), 7.15 – 7.11 (m, 2H), 6.42 (s, 1H), 4.30 (ddd, *J* = 7.7, 5.1, 1.0 Hz, 1H), 4.22 (d, *J* = 5.9 Hz, 2H), 4.17 – 4.10 (m, 1H), 3.64 (t, *J* = 6.4 Hz, 2H), 3.52 – 3.46 (m, 12H), 3.38 (t, *J* = 5.9 Hz, 2H), 3.18 (q, *J* = 5.9 Hz, 2H), 3.09 (ddd, *J* = 8.6, 6.2, 4.4 Hz, 1H), 2.81 (dd, *J* = 12.5, 5.1 Hz, 1H), 2.58 (d, *J* = 12.4 Hz, 1H), 2.43 (d, *J* = 4.7 Hz, 3H), 2.38 (t, *J* = 6.4 Hz, 2H), 2.06 (t, *J* = 7.4 Hz, 2H), 1.61 (ddt, *J* = 13.6, 9.8, 6.1 Hz, 1H), 1.55 – 1.40 (m, 3H), 1.36 – 1.23 (m, 2H); $^{13}$C NMR (151 MHz, DMSO) $\delta_C$ 172.14, 170.0, 162.68, 156.74, 155.35, 153.38, 138.22, 135.92, 133.19, 133.18, 128.94, 127.73, 127.35 (2C), 124.03, 123.63, 119.75 (2C), 105.00, 69.71 (2C), 69.63 (2C), 69.48, 69.47, 69.06, 66.81, 61.00, 59.16, 55.37, 41.59, 39.78, 38.39, 36.09, 35.02, 28.45, 28.11, 27.95, 25.19.

**N-(4-((5-Chloro-4-((2-(N-methylsulfamoyl)phenyl)amino)pyrimidin-2-yl)amino)benzyl)-1-(5-((3aS,4S,6aR)-2-oxohexahydro-1H-thieno[3,4-d]imidazol-4-yl)pentanamido)-3,6,9,12,15,18,21,24-octaoxaheptacosan-27-amide 2,2,2-trifluoroacetate (Cp19-PEG8-biotin):**

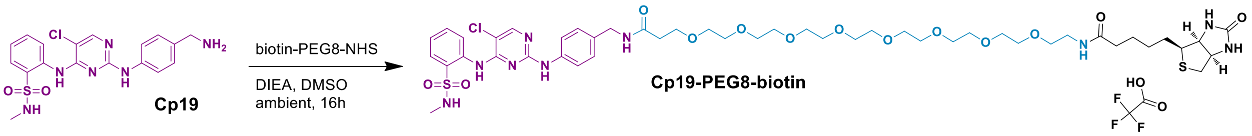


To 2-((2-((4-(Aminomethyl)phenyl)amino)-5-chloropyrimidin-4-yl)amino)-N-methylbenzenesulfonamide (**Cp19**, 10 mg, 0.024 mmol) and biotin-PEG8-NHS (21.9 mg, 0.029 mmol) in 1 mL anhydrous dimethylsulfoxide was added N-ethyl-N-isopropylpropan-2-amine (20 µl, 0.115 mmol) and the reaction shaken at ambient for 16 h. The reaction was diluted to 3 mL with 90% DMSO/water (v/v) and purified by preparative HPLC TFA method 2. Fractions containing the desired peak were combined and lyophilized to give the desired product **Cp19-PEG8-biotin** (12.8 mg, 45.4 %) as a colorless waxy solid: LCMS FA method 1: $R_t$ = 1.35 min, ESI-MS *m/z* 535.2 $(M+2H)^{2+}$, 1068.2 $(M+H)^+$, 1065.6 $(M-H)^-$; LCMS AA method 1: $R_t$ = 1.39 min, ESI-MS *m/z* 535.6 $(M+2H)^{2+}$, 1068.0 $(M+H)^+$, 1066.6 $(M-H)^-$; $^1$H NMR (600 MHz, DMSO) $\delta_H$ 9.57 (s, 1H), 9.39 (s, 1H), 8.50 (d, *J* = 7.7 Hz, 1H), 8.28 (t, *J* = 5.8 Hz, 1H), 8.28 (s, 1H), 7.85 – 7.80 (m, 2H), 7.77 (q, *J* = 4.9 Hz, 1H), 7.65 (ddd, *J* = 8.7, 7.3, 1.6 Hz, 1H), 7.53 (d, *J* = 8.1 Hz, 2H), 7.34 (ddd, *J* = 8.3, 7.4, 1.2 Hz, 1H), 7.16 – 7.11 (m, 2H), 6.41 (s, 2H), 4.31 (ddd, *J* = 7.8, 5.1, 1.0 Hz, 1H), 4.22 (d, *J* = 5.9 Hz, 2H), 4.13 (dd, *J* = 7.7, 4.4 Hz, 1H), 3.64 (t, *J* = 6.4 Hz, 2H), 3.53 – 3.47 (m, 28H), 3.39 (t, *J* = 5.9 Hz, 2H), 3.18 (q, *J* = 5.9 Hz, 2H), 3.09 (ddd, *J* = 8.6, 6.2, 4.4 Hz, 1H), 2.82 (dd, *J* = 12.5, 5.1 Hz, 1H), 2.58 (d, *J* = 12.4 Hz, 1H), 2.44 (d, *J* = 4.8 Hz, 3H), 2.38 (t, *J* = 6.4 Hz, 2H), 2.06 (t, *J* = 7.4 Hz, 2H), 1.61 (ddt, *J* = 13.5, 9.8, 6.1 Hz, 1H), 1.56 – 1.42 (m, 3H), 1.36 – 1.25 (m, 2H); $^{13}$C NMR (151 MHz, DMSO) $\delta_C$ 172.10, 169.97, 162.68, 157.00, 155.20, 153.87, 138.35, 136.00, 133.14, 133.04, 128.93, 127.56, 127.3 (2C), 123.86, 123.52, 119.59 (2C), 104.96, 69.70 (10C), 69.65, 69.62, 69.49, 69.47, 69.09, 66.82, 60.99, 59.16, 55.35, 41.59, 39.77, 38.37, 36.10, 35.01, 28.46, 28.11, 27.95, 25.17.

**N-(4-((5-Chloro-4-((2-(N-methylsulfamoyl)phenyl)amino)pyrimidin-2-yl)amino)benzyl)-1-(5-((3aS,4S,6aR)-2-oxohexahydro-1H-thieno[3,4-d]imidazol-4-yl)pentanamido)-**

**3,6,9,12,15,18,21,24,27,30,33,36-dodecaoxanonatriacontan-39-amide 2,2,2-trifluoroacetate (Cp19-PEG12-biotin):**

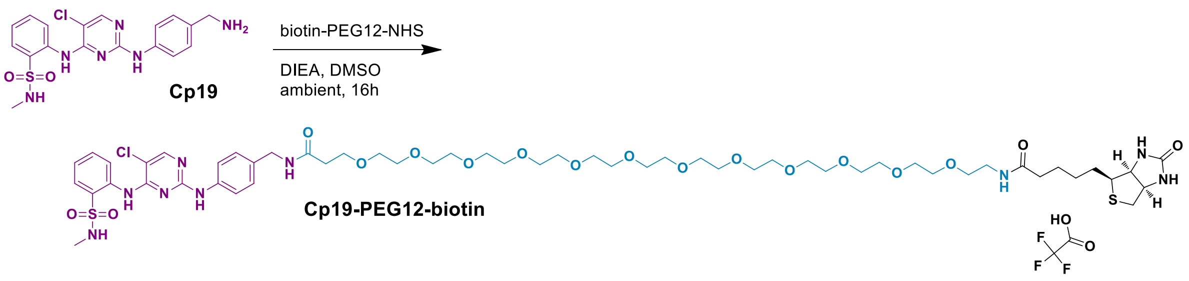


To 2-((2-((4-(Aminomethyl)phenyl)amino)-5-chloropyrimidin-4-yl)amino)-N-methylbenzenesulfonamide (**Cp19**, 10 mg, 0.024 mmol) and biotin-PEG12-NHS (34 mg, 0.036 mmol) in 1 mL anhydrous dimethylsulfoxide was added N-ethyl-N-isopropylpropan-2-amine (20 µl, 0.115 mmol) and the reaction shaken at ambient for 16 h. The reaction was diluted to 3 mL with 90% DMSO/water (v/v) and purified by preparative HPLC TFA method 2. Fractions containing the desired peak were combined and lyophilized to give the desired product **Cp19-PEG12-biotin** (6.3 mg, 19.4 %) as a colorless waxy solid: LCMS FA method 1: $R_t$ = 1.38 min, ESI-MS *m/z* 622.8 $(M+2H)^{2+}$, 1242.8 $(M-H)^-$; LCMS AA method 1: $R_t$ = 1.42 min, ESI-MS *m/z* 622.8 $(M+2H)^{2+}$; $^1$H NMR (600 MHz, DMSO) $\delta_H$ 9.52 (s, 1H), 9.35 (s, 1H), 8.53 (d, *J* = 8.4 Hz, 1H), 8.27 (d, *J* = 4.7 Hz, 2H), 7.82 (ddd, *J* = 8.9, 6.3, 3.5 Hz, 2H), 7.78 (q, *J* = 4.9 Hz, 1H), 7.65 (ddd, *J* = 8.6, 7.3, 1.6 Hz, 1H), 7.55 (d, *J* = 8.2 Hz, 2H), 7.13 (d, *J* = 8.4 Hz, 2H), 6.36 (s, 2H), 4.22 (d, *J* = 5.9 Hz, 2H), 4.13 (dd, *J* = 7.7, 4.4 Hz, 1H), 3.66 – 3.59 (m, 2H), 3.53 – 3.48 (m, 44H), 3.39 (t, *J* = 5.9 Hz, 2H), 3.18 (q, *J* = 5.8 Hz, 2H), 3.09 (ddd, *J* = 8.6, 6.1, 4.4 Hz, 1H), 2.81 (dd, *J* = 12.5, 5.1 Hz, 1H), 2.58 (d, *J* = 12.4 Hz, 1H), 2.44 (d, *J* = 4.8 Hz, 3H), 2.38 (t, *J* = 6.4 Hz, 2H), 2.06 (t, *J* = 7.5 Hz, 2H), 1.66 – 1.57 (m, 1H), 1.55 – 1.42 (m, 3H), 1.36 – 1.25 (m, 2H); $^{13}$C NMR (151 MHz, DMSO) $\delta_C$ 172.10, 169.99, 162.67, 157.32, 155.07, 154.43, 138.55, 136.12, 133.15, 132.90, 128.91, 127.31 (2C), 127.25, 123.66, 123.31, 119.49 (2C), 105.00, 69.75 (20C), 69.58, 69.49, 69.07, 66.84, 61.02, 59.14, 55.38, 41.61, 39.64, 38.39, 36.11, 35.01, 28.46, 28.12, 27.97, 25.19.

**N-(2-(2-(2-((4-((5-chloro-4-((2-(N-methylsulfamoyl)phenyl)amino)pyrimidin-2-yl)amino)benzyl)amino)-2-oxoethoxy)ethoxy)ethyl)-5-((3aS,4S,6aR)-2-oxohexahydro-1H-thieno[3,4-d]imidazol-4-yl)pentanamide-Affi-Gel™10 (Cp19-PEG2-Affi-Gel™10):**

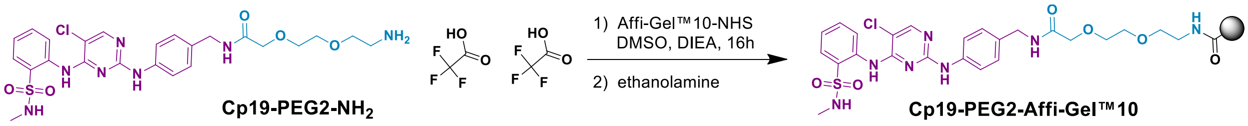


Affi-Gel™10-NHS active ester agarose (12.5 mL, 0.75 mmol) was placed in a 25 mL ISOLUTE™ single-fritted reservoir with a stopcock attached and drained. The resin was washed with 5 × 25 mL anhydrous 2-propanol, 2 × 25 mL anhydrous dimethylsulfoxide. To the washed resin was added a mixture of 2-(2-(2-aminoethoxy)ethoxy)-N-(4-((5-chloro-4-((2-(N-methylsulfamoyl)phenyl)amino)pyrimidin-2-yl)amino)benzyl)acetamide bis-(2,2,2-trifluoroacetate) (**Cp19-PEG2-NH$_2$**, 5.0 mg, 0.00625 mmol) dissolved in 10 mL anhydrous dimethylsulfoxide to which was added N-ethyl-N-isopropylpropan-2-amine (96 µl, 0.562 mmol). The resin container was sealed and gently agitated on a rotary device for 16 h at ambient. The resin was drained, suspended in 10 mL 1 M ethanolamine and shaken for 5 min to quench any remaining active ester. The resin was drained, washed with 2 × 25 mL anhydrous dimethylsulfoxide, 5 × 25 mL anhydrous 2-propanol and stored suspended in 2-propanol at 4 ºC. Loading of the resin and stability of Cp19-PEG2-$NH_2$ through this process was confirmed by testing the original Cp19-PEG2-$NH_2$/DMSO/DIEA mix, 1 M ethanolamine treated Cp19-PEG2-$NH_2$/DMSO/DIEA mix and the first flowthrough after resin loading by LCMS FA method 1. No change in Cp19-PEG2-$NH_2$ retention time or molecular weight were noted except for complete disappearance of the Cp19-PEG2-$NH_2$ peak with resin treatment.

**3-amino-N-(4-(5-amino-6-(1-ethyl-1H-imidazo[4,5-c]pyridin-2-yl)pyrazin-2-yl)phenyl)propanamide 2,2,2-trifluoroacetate (Cp8)** was synthesized by WuXi Apptec, Shanghai, China as described (9)(10).

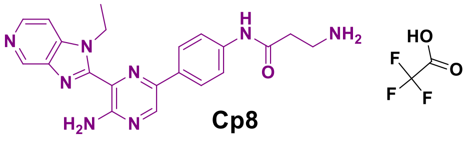


LCMS FA method 1: $R_t$ = 1.24 min, ESI-MS *m/z* 420.0 $(M+H)^+$, 839.6 $(2M+H)^+$; $^1$H NMR (400 MHz, DMSO-$d_6$) $\delta_H$ 1.53 (t, J=7.03 Hz, 3H), 2.75 (t, J=6.79 Hz, 2H), 3.07-3.15 (m, 2H), 5.07 (q, J=6.91 Hz, 2H), 7.72-7.90 (m, 5H), 7.97 (d, J=8.70 Hz, 2H), 8.25 (br s, 2H), 8.36 (d, J=6.44 Hz, 1H), 8.73 (d, J=6.44 Hz, 1H), 8.88 (s, 1H), 9.50 (s, 1H), 10.35 (s, 1H).

(9) Dittus. L., et al, *ACS Chem. Biol.,* **2017**, 12(10), 2515-2521.
(10) Eberl, C.H., et al, *Scientific Reports,* **2019**, 9(1), 1-14.

**N-(3-((4-(5-amino-6-(1-ethyl-1H-imidazo[4,5-c]pyridin-2-yl)pyrazin-2-yl)phenyl)amino)-3-oxopropyl)-1-(5-((3aS,4S,6aR)-2-oxohexahydro-1H-thieno[3,4-d]imidazol-4-yl)pentanamido)-3,6,9,12,15,18,21,24-octaoxaheptacosan-27-amide 2,2,2-trifluoroacetate (Cp8-PEG8-biotin):**

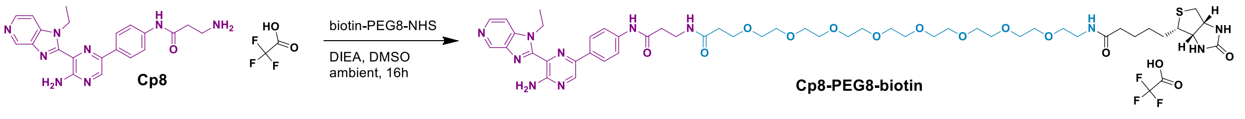


To 3-amino-N-(4-(5-amino-6-(1-ethyl-1H-imidazo[4,5-c]pyridin-2-yl)pyrazin-2-yl)phenyl)propanamide 2,2,2-trifluoroacetate (**Cp8**, 22 mg, 0.043 mmol) and biotin-PEG8-NHS (32.6 mg, 0.043 mmol) in 1 mL anhydrous dimethylsulfoxide was added N-ethyl-N-isopropylpropan-2-amine (20 µl, 0.115 mmol) and the reaction shaken at ambient for 16 h. The reaction was diluted to 3 mL with 90% DMSO/water (v/v) and purified by preparative HPLC TFA method 2. Fractions containing the desired peak were combined and lyophilized to give the desired product **Cp8-PEG8-biotin** (29.3 mg, 59.0 %) as a waxy solid: LCMS FA method 1: $R_t$ = 1.07 min, ESI-MS *m/z* 351.4 $(M+3H)^{3+}$, 526.6 $(M+2H)^{2+}$, 1051.6 $(M+H)^+$. LCMS AA method 1: $R_t$ = 1.32 min, ESI-MS *m/z* 351.2 $(M+3H)^{3+}$, 527.0 $(M+2H)^{2+}$, 1051.6 $(M+H)^+$. $^1$H NMR (500 MHz, DMSO) δ 10.10 (s, 1H), 9.51 (s, 1H), 8.89 (s, 1H), 8.73 (d, *J* = 6.5 Hz, 1H), 8.38 (d, *J* = 6.5 Hz, 1H), 8.23 (s, 1H), 7.97 (dd, *J* = 13.9, 7.2 Hz, 3H), 7.81 (t, *J* = 5.6 Hz, 1H), 7.77 (d, *J* = 8.8 Hz, 2H), 6.40 (s, 1H), 6.35 (s, 1H), 5.08 (q, *J* = 7.0 Hz, 2H), 4.30 (dd, *J* = 7.8, 5.0 Hz, 1H), 4.18 – 4.06 (m, 1H), 3.59 (t, *J* = 6.5 Hz, 2H), 3.52 – 3.44 (m, 34H), 3.17 (q, *J* = 6.0 Hz, 3H), 3.08 (dt, *J* = 8.9, 5.5 Hz, 1H), 2.82 – 2.73 (m, 1H),

2.32 (t, $J$ = 6.4 Hz, 2H), 2.05 (t, $J$ = 7.3 Hz, 2H), 1.63 – 1.57 (m, 2H), 1.54 (t, $J$ = 7.0 Hz, 3H), 1.47 (dd, $J$ = 8.9, 5.8 Hz, 3H), 1.36 – 1.20 (m, 2H).

**6-Acetyl-8-cyclopentyl-5-methyl-2-((5-(piperazin-1-yl)pyridin-2-yl)amino)pyrido[2,3-d]pyrimidin-7(8H)-one bis-(hydrochloric acid) (Palbociclib)** was synthesized by Pharmaron, Beijing, China as described (11).

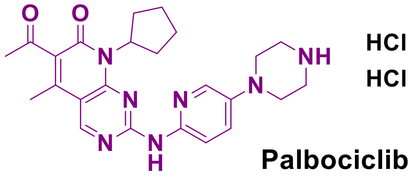


LCMS FA method 1: $R_t$ = 0.81 min, ESI-MS *m/z* 224.6 $(M+2H)^{2+}$, 448.0 $(M+H)^+$; $^1$H NMR (300 MHz, DMSO-d6) $\delta_H$ 0.81 - 0.98 (m, 2H), 1.11 - 1.36 (m, 7H), 1.37 - 1.50 (m, 10H), 1.52 - 1.67 (m, 2H), 1.69 - 1.82 (m, 2H), 1.84 - 1.96 (m, 2H), 1.97 - 2.07 (m, 2H), 2.11 - 2.31 (m, 2H), 2.33 - 2.45 (m, 3H), 3.06 - 3.20 (m, 4H), 3.43 - 3.61 (m, 4H), 3.76 - 3.90 (m, 2H), 3.97 - 4.12 (m, 2H), 4.40 - 4.55 (m, 1H), 5.74 - 5.94 (p, J = 8.6 Hz, 1H), 7.42 - 7.58 (dd, J = 3.0, 8.9 Hz, 1H), 7.85 - 7.98 (d, J = 9.0 Hz, 1H), 8.01 - 8.17 (d, J = 2.9 Hz, 1H), 8.81 - 8.95 (s, 1H), 9.92 - 10.10 (s, 1H).

(11) Toogood, P.L., et al., *J. Med. Chem.*, **2005**, *48(7),* 2388-2406.

**6-Acetyl-2-((5-(4-(3-aminopropyl)piperazin-1-yl)pyridin-2-yl)amino)-8-cyclopentyl-5-methylpyrido[2,3-d]pyrimidin-7(8H)-one bis-(2,2,2-trifluoroacetate) (Palbociclib-2):**

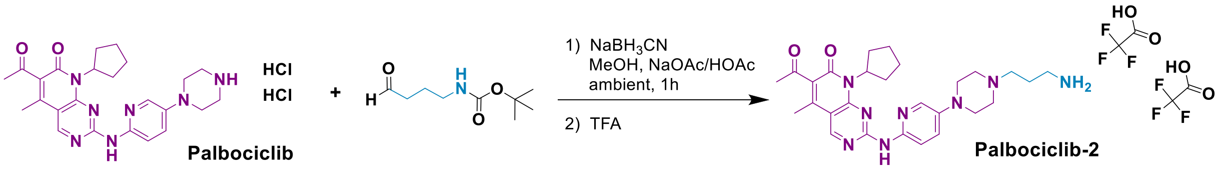


To 6-Acetyl-8-cyclopentyl-5-methyl-2-((5-(piperazin-1-yl)pyridin-2-yl)amino)pyrido[2,3-d]pyrimidin-7(8H)-one bis-(hydrochloric acid) (**Palbociclib**, 93.2 mg, 0.179 mmol) and *tert*-butyl (3-oxopropyl)carbamate (37.2 mg, 0.215 mmol) dissolved in 2 mL MeOH pH 5 buffer (containing 24 g sodium acetate, 18 g acetic acid in 500 mL methanol) was added sodium cyanoborohydride (24.8 mg, 0.394 mmol) and the reaction shaken at ambient for 1 h. The reaction was diluted with 20 mL water and extracted with 5 x 20 mL dichloromethane. The combined organic layers were dried over magnesium sulfate and evaporated to give the Boc protected intermediate as a yellow solid that was carried forward crude due to solubility issues (124.3 mg, 115 %); LCMS FA method 1: $R_t$ = 1.09 min, ESI-MS *m/z* 303.0 $(M+2H)^{2+}$, 605.0 $(M+H)^+$.

**Palbociclib-2-Boc** (impure, 124.3 mg) was dissolved in 1 mL trifluoroacetic acid and immediately evaporated to dryness under a stream of dry nitrogen gas. The residue was dissolved to 3 mL with 90% DMSO/water (v/v) and purified by preparative HPLC TFA method 2. Fractions containing the desired peak were combined and lyophilized to give the desired product **Palbociclib-2** (62.8 mg, 47.8 % over two steps) as a yellow solid: LCMS FA method 1: $R_t$ = 0.68 min, ESI-MS *m/z* 253.0 $(M+2H)^{2+}$, 505.2 $(M+H)^+$. LCMS

AA method 1: $R_t$ = 1.47 min, ESI-MS *m/z* 505.2 $(M+H)^+$. $^1$H NMR (500 MHz, DMSO) δ 9.73 (s, 1H), 8.92 (s, 1H), 8.19 – 7.98 (m, 1H), 7.98 – 7.81 (m, 4H), 7.52 (dd, *J* = 9.1, 3.1 Hz, 1H), 5.81 (p, *J* = 8.8 Hz, 1H), 3.76 (s, 3H), 3.46 (s, 2H), 3.31 (s, 2H), 3.19 (d, *J* = 2.3 Hz, 2H), 2.94 (t, *J* = 7.4 Hz, 2H), 2.42 (s, 3H), 2.32 (s, 5H), 2.11 – 1.99 (m, 2H), 1.98 – 1.86 (m, 2H), 1.79 (dddt, *J* = 11.5, 8.8, 5.8, 3.6 Hz, 2H), 1.68 – 1.50 (m, 2H).

**N-(3-(4-(6-((6-acetyl-8-cyclopentyl-5-methyl-7-oxo-7,8-dihydropyrido[2,3-d]pyrimidin-2-yl)amino)pyridin-3-yl)piperazin-1-yl)propyl)-1-(5-((3aS,4S,6aR)-2-oxohexahydro-1H-thieno[3,4-d]imidazol-4-yl)pentanamido)-3,6,9,12,15,18,21,24-octaoxaheptacosan-27-amide bis-(2,2,2-trifluoroacetate) (Palbociclib-2-PEG8-biotin):**

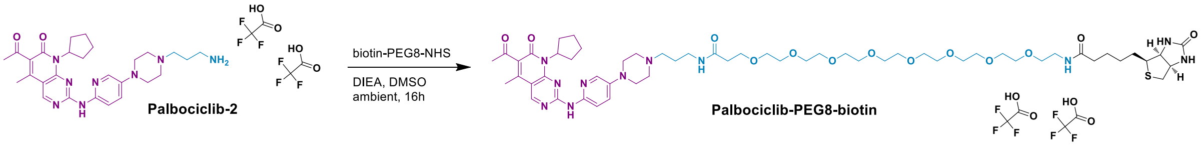


To 6-acetyl-2-((5-(4-(3-aminopropyl)piperazin-1-yl)pyridin-2-yl)amino)-8-cyclopentyl-5-methylpyrido[2,3-d]pyrimidin-7(8H)-one bis-(2,2,2-trifluoroacetate) (**Palbociclib-2**, 24.2 mg, 33 mmol)

and biotin-PEG8-NHS (30.3 mg, 0.040 mmol) in 1 mL anhydrous dimethylsulfoxide was added N-ethyl-N-isopropylpropan-2-amine (23 µl, 0.132 mmol) and the reaction shaken at ambient for 16 h. The reaction was diluted to 3 mL with 90% DMSO/water (v/v) and purified by preparative HPLC TFA method 2. Fractions containing the desired peak were combined and lyophilized to give the desired product **Palbociclib-2-PEG8-biotin** (23.1 mg, 50.5 %) as a waxy solid: LCMS FA method 1: $R_t$ = 1.04 min, ESI-MS *m/z* 385.2 $(M+3H)^{3+}$, 577.6 $(M+2H)^{2+}$. LCMS AA method 1: $R_t$ = 1.39 min, ESI-MS *m/z* 385.6 $(M+3H)^{3+}$, 577.8 $(M+2H)^{2+}$, 1155.8 $(M+H)^+$. $^1$H NMR (600 MHz, DMSO) δ 10.25 (s, 1H), 9.64 (s, 1H), 8.97 (s, 1H), 8.13 (d, *J* = 3.0 Hz, 1H), 8.08 (t, *J* = 5.9 Hz, 1H), 7.92 (d, *J* = 9.0 Hz, 1H), 7.81 (t, *J* = 5.7 Hz, 1H), 7.59 (dd, *J* = 9.1, 3.1 Hz, 1H), 6.40 (s, 1H), 6.35 (s, 1H), 5.83 (p, *J* = 8.9 Hz, 1H), 4.30 (dd, *J* = 7.7, 5.0 Hz, 1H), 4.13 (dd, *J* = 7.8, 4.5 Hz, 1H), 3.88 (d, *J* = 13.0 Hz, 2H), 3.66 – 3.56 (m, 44H), 2.82 (dd, *J* = 12.4, 5.1 Hz, 1H), 2.58 (d, *J* = 12.4 Hz, 1H), 2.43 (s, 3H), 2.35 (t, *J* = 6.4 Hz, 2H), 2.32 (s, 3H), 2.25 (dd, *J* = 11.8, 7.8 Hz, 2H), 2.06 (t, *J* = 7.4 Hz, 2H), 1.94 – 1.72 (m, 7H), 1.66 – 1.56 (m, 3H), 1.51 – 1.37 (m, 2H), 1.36 – 1.15 (m, 2H).
